# Action-value processing underlies the role of the dorsal anterior cingulate cortex in performance monitoring during self-regulation of affect

**DOI:** 10.1101/2020.09.08.283671

**Authors:** Keith A. Bush, G. Andrew James, Anthony A. Privratsky, Kevin P. Fialkowski, Clinton D. Kilts

## Abstract

In this study, we merged methods from engineering control theory, machine learning, and human neuroimaging to critically test the putative role of the dorsal anterior cingulate cortex (dACC) in goal-directed performance monitoring during an emotion regulation task. Healthy adult participants (n=94) underwent cued-recall and re-experiencing of their responses to affective image stimuli with concurrent functional magnetic resonance imaging and psychophysiological response recording. During cued-recall/re-experiencing trials, participants engaged in explicit self-regulation of their momentary affective state to match a pre-defined affective goal state. Within these trials, neural decoding methods measured affect processing from fMRI BOLD signals across the orthogonal affective dimensions of valence and arousal. Participants’ affective brain states were independently validated via facial electromyography (valence) and electrodermal activity (arousal) responses. The decoded affective states were then used to contrast four computational models of performance monitoring (i.e., error, predicted response outcome, action-value, and conflict) by their relative abilities to explain emotion regulation task-related dACC activation. We found that the dACC most plausibly encodes action-value for both valence and arousal processing. We also confirmed that dACC activation directly encodes affective arousal and also likely encodes recruitment of attention and regulation resources. Beyond its contribution to improving our understanding of the roles that the dACC plays in emotion regulation, this study introduced a novel analytical framework through which affect processing and regulation may be functionally dissociated, thereby permitting mechanistic analysis of real-world emotion regulation strategies, e.g., distraction and reappraisal, which are widely employed in cognitive behavioral therapy to address clinical deficits in emotion regulation.

## Introduction

The expression and perception of emotions are valuable social cognitive resources that allow us to focus our attention to salient environmental features^1^, orchestrate social exchanges^2^, prioritize our decisions^3^, and engage in appetitive and aversive behaviors^4^. Conversely, disruption of the multiple roles that emotions perform in directing our cognitions and behaviors is associated with mental health problems. Emotion misprocessing (e.g., emotional hypo/hyperreactivity and lability) engenders maladaptive emotional states implicated in psychiatric illnesses^5^. Thus, we routinely deploy emotion regulation strategies to mitigate such processing errors and to mold our emotional experiences in ways that benefit our well-being^6^. Critical to the success of such regulation strategies is the ability to obtain feedback about momentary states of affect relative to an affective goal state, termed performance monitoring. The present study sought to utilize computational approaches to compare and contrast the roles of multiple performance monitoring signals theoretically related to the regulation of emotion.

The process model of emotion regulation^7,8^ provides a conceptual bridge between the temporal steps comprising emotion formation (i.e., the modal model^9^) and the multitude of cognitive processes serving to extinguish, alter, or promote such formation. For example, two widely deployed emotion regulation strategies, distraction and reappraisal, respectively target attentional deployment versus cognitive appraisal components of the process model in order to regulate the prepotent (and potentially dysfunctional) trajectory of emotional response. Extant empirical studies of emotion regulation in healthy subjects describe the comparative efficacy of these strategies^10^, individual differences in strategy efficacy^11^, as well as their attendant functional neurocircuitry^12–14^. It is the process model that provides an overarching framework by which to organize these studies and to extrapolate their findings to inform observed patterns of emotion dysregulation associated with psychopathology^15–19^.

Computational models of emotion regulation, therefore, should build on the process model’s components to elaborate descriptions of its underlying mechanisms^20^ by merging empirical observations with mathematical rigor. A heuristic framework for model development is provided by the superordinate process of cognitive control, which has long drawn mechanistic inference from engineering control theory^21–23^ to empirically test computational models^24,25^ of its mechanisms of control. As a step in this direction, leading scholars recently proposed a neuroanatomically-constrained unified emotion regulation framework^26^ rooted in reinforcement learning^27^, a value-based multi-step decision strategy having important mathematical and conceptual connections to dynamic programming^28^, optimal control theory^29^, and machine learning^30^ with theorized links to cognitive control^21^.

Building on these ideas, this study formally tested the computational basis of performance monitoring signals^31^ during emotion regulation. To accomplish this goal, we constrained our study’s focus to the top-down cognitive control of affect processing^20,32^ (i.e., explicit regulation^33^) and, drawing on results from prior work within the cognitive control literature, assigned patterns of functional neuroanatomical activation associated with explicit affect regulation to the roles performed by subcomponents of a formal engineering control system. These subcomponents included the plant (the system to be controlled, i.e., affect processing) and the control law (the system that monitors control performance and acts to best align the plant’s state with a desired goal state). To approximate a relevant control law we input extant theoretical measures of performance monitoring, previously posited by the cognitive control and emotion regulation research communities, into this computational model and measured the concordance between the model’s predictions and neural activation patterns observed in the dorsal anterior cingulate cortex (dACC), a region consistently linked to performance monitoring^24,31,34–46^. Thus, this study sought to merge the potential of computational neuroscience with the robust field of engineering control theory to provide a fresh perspective on the role of the dACC in emotion regulation.

## Methods

### Study Overview

This study analyzed behavioral, psychophysiological, and functional brain imaging data acquired from two separate experiments, the Intrinsic Neuromodulation of Core Affect (INCA) experiment and the Cognitive Control Theoretic Mechanisms of Real-time fMRI-Guided Neuromodulation (CTM) experiment (National Science Foundation, BCS-1735820). Both experiments were based in control theoretic functional neuroimaging explorations of the brain representations of affect processing and regulation, and incorporated both unguided and real-time fMRI-guided affect regulation tasks. Importantly, both experiments shared identical affective stimuli as well as identical design and ordering of the affect processing, affect regulation, and resting state tasks. The following expands on the experimental design details that enabled the present affect regulation analysis.

We conducted both the INCA and CTM studies over two separate sessions, each occurring on separate days. During Session 1, participants provided written informed consent, received screening for clinically relevant exclusionary criteria via structured clinical interview, and completed behavioral assessments and questionnaires. We acquired magnetic resonance imaging (MRI) and concurrent psychophysiological measurements during Session 2. The analyses reported here focus on data acquired during the first three functional image acquisitions (i.e., scans) of Session 2, which correspond to the two System Identification task scans and one Resting State task scan. The relevant task descriptions are elaborated below.

### Experimental Task Design and Conceptual Model

The System Identification task performed two independent roles in our study. First, this task implicitly induced, via visual image stimuli, *affect processing* in our participants that was measured via concurrent functional MRI (fMRI) and psychophysiology in order to construct and validate study-specific neural decoding models of affect processing. Second, this task, via cued-recall and re-experiencing trials, induced explicit *affect regulation* to attain an objectively known goal, allowing for the measurement of moment-to-moment affect regulation performance. By this design, isolation of the specific process of explicit affect regulation did not rely on the perhaps more conventional use of affective state manipulation tasks such as re-appraisal or distraction, but rather focused on the effortful re-creation of a prior implicitly induced affective state (goal state) to best approximate the relevant components of the engineering control system framework.

We induced the affect processing state using 90 image stimuli that were computationally sampled from the International Affective Picture Set (IAPS) to maximize the range of valence and arousal processing demands induced by the resultant set of image stimuli^47,48^ (see Supplemental Figure S1). We presented each image stimulus for 2 s followed by an inter-trial interval (ITI) uniformly randomly sampled from the range 2–6 s during which we presented a fixation cross. We labeled these image presentation sequences as implicit affect processing induction trials.

We induced the affect regulation goal state using 30 image stimuli (independently but identically sampled as described above from the remaining IAPS images, see Supp. Fig. S1) as part of the experiment’s cued-recall/re-experiencing trials (see Figure 1, panel A). In these trials, we presented a cue image stimulus for 2 s followed by a visual preparation instruction (the word “FEEL” superimposed over the still-observed image) for 2 s followed by a recall/re-experiencing instruction in which the image disappears (leaving only the word “FEEL”) for 8 s. Participants were instructed to respond to the “FEEL” prompt during recall/re-experiencing trials by explicitly regulating their affect processing state to match the affect processing state induced by implicit responses to the cue image stimulus. Finally, a fixation cross replaced the word “FEEL” for an ITI sampled uniformly randomly from the range 2–6 s. We thus induced participants to regulate their affect state relative to a goal state, upon seeing the recall/re-experience instruction (i.e., “FEEL”), according to the following instructions. “[W]hen the image disappears and just the word ‘feel’ remains, we want you to re-imagine the image you just saw and try to re-feel how the image made you feel when you first saw it. Hold that feeling the entire time the word ‘feel’ is on the screen.” The complete list of IAPS images used in this experiment, including their task role and normative valence and arousal properties, are summarized in Supplemental Table S1.

**Figure 1:**
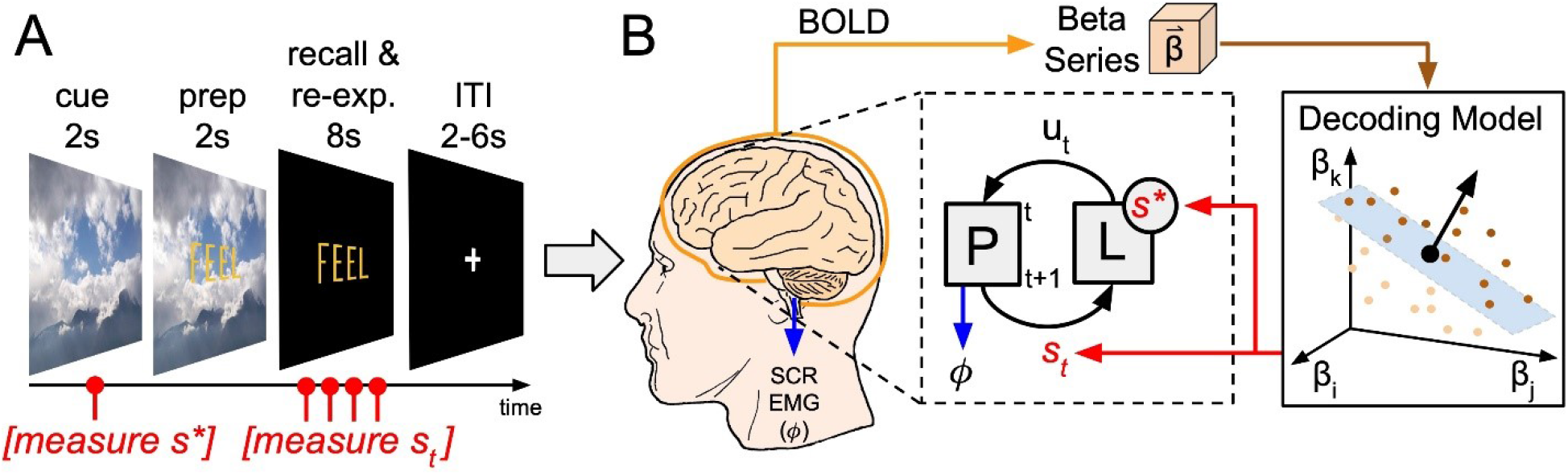
Explicit affect regulation task design and conceptual model of affect regulation in an engineering control system framework. **(A)** Schematic of a cued-recall/re-experiencing trial. The trial presents a cue image stimulus for 2 s. The trial then overlays the word “FEEL” for 2 s in yellow font. The image disappears, replaced by a black screen, leaving only the word “FEEL” for 8 s. The trial ends with a fixation cross for an inter-trial interval sampled uniformly randomly on the range of 2-6 s. **(B)** A closed-loop conceptual model of the brain processing related to explicit affect regulation induced by the cued-recall/re-experiencing task. The proposed closed-loop control system’s box diagram falls within the dashed area of the figure. The system of interest, aka the plant, P, is assumed to be observable according to measurements of its state, s. In this study, the plant is composed of the independent dimensions of affect processing, valence and arousal. The control law, L, evaluates the plant’s momentary state (i.e., performance monitoring) and responds with a control action, u, intended to perturb P such that the system state moves closer to the desired state, i.e., the goal, s*. Each cued-recall/re-experiencing task trial first captures the goal state and then captures four separate dynamic measurements (brain state representations and psychophysiological responses) of the system state as the subject explicitly regulates their affective state to match the goal state. Brain representations of the goal and system states are measured by submitting whole-brain gray matter patterns of neural activation extracted from the subject’s fMRI BOLD signal to neural decoding models of affective valence and arousal processing. To characterize s* and the momentary states of P, psychophysiological measures of arousal (skin conductance response, SCR) and valence (facial electromyography, EMG) are captured concurrently with fMRI BOLD signal.

Conceptually, our System Identification task facilitated the construction of two control system subcomponents necessary to conduct our proposed test of dACC function. Implicit affect processing induction trials yielded fMRI-derived brain states from which we built neural decoding models of affect processing. These decoding models (see Fig. 1, panel B) served as the computational representation of the state of the plant, allowing us to quantitatively measure affect processing during regulation. The cued-recall/re-experiencing trials provided brief tests of the processes of explicit affect regulation. Critically, the cue conditions of these trials induced subject-specific implicit affect processing that was captured by the decoding model, forming objective goal states for post-hoc analysis of subsequent regulation performance. The trajectory of affect processing (our measure of affect regulation relative to the goal state) was decoded post-hoc from the sequence of fMRI-derived brain states induced by the recall/re-experiencing condition’s instruction (“FEEL”) to explicitly regulate one’s affect to achieve and maintain the goal state. Theoretical measures of performance monitoring, which we hypothesize are executed by the control law within the dACC, were then constructed from these affect processing measurements (see Fig. 1, panel B).

Finally, we acquired 7.5 min of fMRI resting state data in which subjects performed mind-wandering. During training, subjects were instructed to “let your mind wander, not focusing on any specific thought” and to “try to keep your head still and your eyes open [and] … blink naturally.” We used resting state fMRI as a control condition by which to measure baseline fluctuations of affect processing dynamics in the untasked brain.

### Ethics Statement

All participants provided written informed consent after receiving written and verbal descriptions of the study procedures, risks, and benefits. We performed all study procedures and data analysis with approval and oversight of the Institutional Review Board at the University of Arkansas for Medical Sciences (UAMS) in accordance with the Declaration of Helsinki and relevant institutional guidelines and policies.

### Participants

We sought to enroll study participants who lived within a one hour drive of the UAMS campus in Little Rock, Arkansas who responded to recruitment materials in the forms of publicly posted flyers, direct emails to identified research volunteers in the ARresearch registry (arresearch.org), and social media advertisements. From our total participant sample (n=97), we excluded from analysis two subjects enrolled in the CTM study who did not complete the resting state task scan (due to early exit from the scanner) as well as one subject enrolled in the INCA study who was inadvertently included in the study despite meeting exclusionary criteria. The final participant sample (n=94; n_CTM_=75 and n_INCA_=19) possessed the following demographic characteristics: age [mean(s.d.)]: 36.6(13.8), range 18-64; sex: 61(65%) female, race/ethnicity: 80(85.1%) self-reporting as White or Caucasian, 11(11.7%) as Black or African-American, 1(1.1%) as Asian, and 2 (2.1%) reporting as more than one race; education [mean(s.d.)]: 16.7(2.6) years, range 12-23; WAIS-IV IQ [mean(s.d.)]: 105.8(14.0), range 74-145. All of the study’s participants were right-handed native-born United States citizens who were medically healthy and exhibited no current Axis I psychopathology, including mood disorders, as assessed by the SCID-IV clinical interview^49^. All participants reported no current use of psychotropic medication and produced a negative urine screen for drugs of abuse (cocaine, amphetamines, methamphetamines, marijuana, opiates, and benzodiazepines) immediately prior to the MRI scan. CTM participants also produced a negative urine screen prior to SCID-IV clinical interview. When necessary, we corrected participants’ vision to 20/20 using an MRI compatible lens system (MediGoggles™, Oxfordshire, United Kingdom), and we excluded all participants endorsing color blindness.

### MR Image Acquisition and Preprocessing

We acquired all imaging data for the INCA and CTM experiments using the same Philips 3T Achieva X-series MRI scanner (Philips Healthcare, Eindhoven, The Netherlands) with a 32-channel head coil. We acquired anatomic images using an MPRAGE sequence (matrix = 256 × 256, 220 sagittal slices, TR/TE/FA = 8.0844/3.7010/8°, final resolution =0.94 × 0.94 × 1 mm^3^). We acquired functional images using the following EPI sequence parameters: TR/TE/FA = 2000 ms/30 ms/90°, FOV = 240 × 240 mm, matrix = 80 × 80, 37 oblique slices, ascending sequential slice acquisition, slice thickness = 2.5 mm with 0.5 mm gap, final resolution 3.0 × 3.0 × 3.0 mm^3^. We performed all MRI preprocessing using AFNI^50^ (Version AFNI_19.1.04) unless otherwise noted. We processed anatomical data according to the following sequence of steps: skull stripping, spatial normalization to the MNI152 brain atlas, and segmentation (via FSL^51^) into white matter (WM), gray matter (GM), and cerebrospinal fluid (CSF). From the individual participant GM segmentations we constructed a group-level GM mask composed of voxels in which ≥ 50% of individuals exhibited the presence of GM. Our pipeline processed functional images according to the following sequence of steps: despiking, slice-time correction, deobliquing, motion correction, transformation to the spatially normalized anatomic image, regression of the mean time courses and temporal derivatives of the WM and CSF masks as well as a 24-parameter motion model^52,53^, spatial smoothing (8 mm FWHM Gaussian kernel), and scaling to percent signal change. For resting state functional images we performed global mean signal subtraction prior to smoothing and scaling.

### Psychophysiology Data Acquisition and Preprocessing

We acquired putative psychophysiological correlates of affect processing using the BIOPAC MP150 Data Acquisition System (BIOPAC Systems, Inc., Goleta, CA) in conjunction with AcqKnowledge software. We simultaneously captured multiple physiological modalities: galvanic skin response (EDA 100C-MRI module), pulse plethysmography (TSD200-MRI module), respiration transduction (TSD221-MRI module), and facial electromyography (EMG100C-MRI module). We acquired galvanic skin responses from electrodes placed on the medial portions of the thenar and hypothenar eminences of the left hand ^48^. We also captured two separate measurements of facial electromyography (EMG), zygomaticus major (zEMG) and corrugator supercilii (cEMG) responses, using the electrode placement guidelines reported in Fridlund and Cacciopo (1986)^54^. Physiological signals were acquired at 2000 Hz.

We used the canonical skin conductance response function^55^ in conjunction with the beta-series method^56^ to capture temporally succinct physiological correlates of autonomic arousal associated with affect processing induced by both the implicit induction trials^57^ and cued-recall/re-experiencing task trials. In cue-recall/re-experiencing task trials we extracted arousal responses for the cue condition to define the subsequent goal state, s*, as well as at 2 s intervals commencing with the onset of the “FEEL”-prompted recall/re-experiencing condition (see Fig. 1, panel A) to define the momentary states, s, of P. Similarly, we modeled facial electromyography as an independent physiological measure of hedonic valence. We first bandpass filtered the raw EMG signal on the range 10–500 Hz to remove artifacts^58^, rectified the filtered signal, and then extracted the sum of the rectified signal for each relevant task period as our feature set, e.g., over the 2 s cue stimulus as well as over 2 s intervals (commencing at condition onset) of the recall/re-experiencing condition (4 intervals total, see Fig. 1, panel A). Note, due to changes in neuroimaging procedures put in place in response to the COVID-19 pandemic, the final 20 participants of the CTM study were required to wear surgical masks at all times up to insertion as well as removal from the MRI bore, making zEMG placement infeasible. Therefore, zEMG was not acquired from these subjects.

### Decoding Affect Processing

In this work we relied on neural decoding models that have been extensively tested and validated in prior work^47,48,59,60^. We detail key aspects of the modeling processing as follows.

#### Feature and Label Extraction

We combined the canonical hemodynamic response function and the beta-series method^56^ to capture whole-brain gray matter patterns of neural activation in response to the implicit affect processing induction trials within the System Identification task fMRI scans. We refer to these patterns as affective brain states. We then paired these states with class assignment labels {+1,1} derived from the relationship of the known IAPS normative valence and arousal scores (see Supplemental Tables S2 & S3) of the corresponding image stimuli to the middle Likert IAPS score (5 of a 9-point scale). These brain states and class labels form the set of feature-label pairs (one pair each for the independent dimensions of valence and arousal). Also, for validation purposes, we created a second set of features for each subject by projecting the whole-brain gray matter patterns of neural activation to 90-dimensional orthogonal features according to the Gram-Schmidt process^61^.

#### Within-Subject Decoding of Affect Processing

As in earlier work^47,48^, we fit neural decoding models of affect processing according to a support vector machine (SVM) model (using Matlab’s fitcsvm function and default hyperparameters). We modeled affect processing within each subject separately for valence and arousal. These neural decoding models represent the plant of each subject’s control system. We estimated decoding model performance accuracy based on within-subject leave-one-out-cross-validation (LOOCV). For each hold-out feature-label pair (i.e, the test set of the LOOCV), we randomly balanced the training dataset to insure a null accuracy of 0.5 prior to model fitting. For each test set, we repeated this sampling and model fitting process 30 times. The mean prediction accuracy over these samples formed the accuracy of each test set. We then calculated the mean accuracy over all test sets to estimate the prediction accuracy for a single subject.

In contrast to prior work, however, here we took the additional step of re-executing all within-subject cross-validations in a manner where the training dataset’s class-labels were uniformly randomly assigned to features in order to establish the true null distribution of our fitting process according to permutation testing^62^. We then compared the group-level accuracy (mean accuracy across all within-subject models) against the permutation-derived group-level null accuracy in order to report statistical significance of our prediction accuracies. We also substituted the within-subject permutation-derived accuracy as the null probability of the binomial distribution in order to report statistical significance of single-subject decoding performance.

#### Decoding Performance Validation

We designed our neural decoding methodology to be able to generalize to novel task environments (e.g., affect regulation) due to the unknown valence and arousal processing demands these tasks induce. In particular, the IAPS images we used to induce affect processing for training of the neural decoding models were computationally selected^48^ to maximize the difficulty of the classification problem by sampling the full range of valence-arousal experiences available in the IAPS stimuli set (see Supp. Fig. S1), including weakly valent and neutrally arousing experiences. We have shown in prior work^48^ that our affect processing decoding performance is equivalent to the best available performance in the literature by a method that identifies and measures decoding performance of reliable stimuli (i.e, those stimuli that induce consistent affective experiences across subjects). Indeed, reliable stimuli cluster at the affective extremes similar to hand-generated IAPS datasets used in prior work to produce group-level significant and within-subject significant classification accuracies^63^. We executed the reliable stimuli sampling method in this work (see Supplemental Methods) to allow comparison of our decoding performance with the broader affective neuroscience literature.

### Decoding Affect Regulation

Recent work supports out-of-sample application of affect processing neural decoding models^59,64–66^. We decoded affect processing within cued-recall/re-experiencing trials by extracting two separate beta-series per trial: (1) the cue presentation condition (see Fig. 1, panel A), indexed by i, which yielded series β(i); and (2) the four individually acquired volumes related to the recall/re-experiencing condition following the cue condition of each trial (see also Fig. 1, panel A), each volume indexed by t given i, which yielded series β(t|i), where t|i referred to the t^th^ recall/re-experiencing volume succeeding the i^th^ cue. We then decoded the resultant beta-series, yielding SVM hyperplane classification distances, 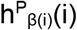 and 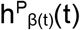, for all i and t where: (1) P denoted the plant of interest, P∈{valence,arousal}, (2) subscripted β(·) denoted the index of the beta-series feature input to the decoding model, and (3) (·) denoted the index of the decoding. In the base case, described here, these indices matched, but they differed, e.g., when we applied the model to forecast future affect processing states. As a final step, we applied Platt scaling^67^ to convert our models’ predictions of hyperplane distance (i.e., distance from a classification decision boundary) to continuous predictions of the affective state’s probability of conforming to the positive class (i.e., the probability of positive valence or high arousal) such that

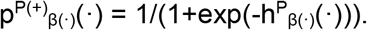

We validated this transformation of our model predictions according to two tests. First, we applied Platt scaling to the decodings (and normative scores) of the implicit affect induction trials and tested, via a linear mixed-effects model, the group-level significance of the models’ predictive fidelity in the transformed probability space. Second, we repeated this validation test for the cue condition stimuli of the cued-recall/re-experiencing task trials using their IAPS normative scores as labels.

### Cued-Recall/Re-experiencing Task Control Condition Modeling

Attempts to identify the brain representations of explicit affect regulation rely on establishing a counter factual condition reflecting the brain states related to affect state when it is not explicitly regulated. In previous work we demonstrated (and validated via independent psychophysiological measures of affect state) the application of neural decoding models to reveal moment-to-moment affect processing dynamics entrained in the resting-state fMRI BOLD signal^59^. In brief, for each subject and for each time point, this approach averaged the neurally decoded affect processing of beta-series constructed from sets of uniformly randomly sampled onset times. Here we leveraged this methodology to construct surrogate cued-recall/re-experiencing tasks within resting state task data which provided experimental controls representing non-explicitly regulated affect states against which to measure the significance of affect processing dynamics entrained during explicit affect regulation. From each subject’s resting state task fMRI data we constructed affect processing estimates, respectively, for valence and arousal. We then uniformly randomly sampled (n=30) onset times of cue stimuli and extracted the attendant cue and recall/re-experiencing affect processing predictions according to the trial’s structure (see Fig. 1, panel A).

### Computational Models of Performance Monitoring

We designed the cued-recall/re-experiencing task as a means of exploring the mathematical structure of regulation process-related performance monitoring to define the functional attributes of dACC activation during explicit affect regulation. From the cognitive control research literature, we initially identified five influential explanatory models of dACC activation (see Table 1); however, upon inspection, we restricted our exploration to just those three models which most closely align with the performance monitoring function of the control law in the engineering control literature and would be best evaluated by our experiment design. We included error processing^68^, which directly represents the plant’s current state with respect to a goal (closed-loop control), predicted response outcome^25^, which estimates the plant’s likely future state with respect to a goal (model-based control), and expected value of control^21^, which estimates the expected value (with respect to a goal, operationalized as a reward function) of making a particular control decision in the current state (reinforcement learning).

**Table 1:**
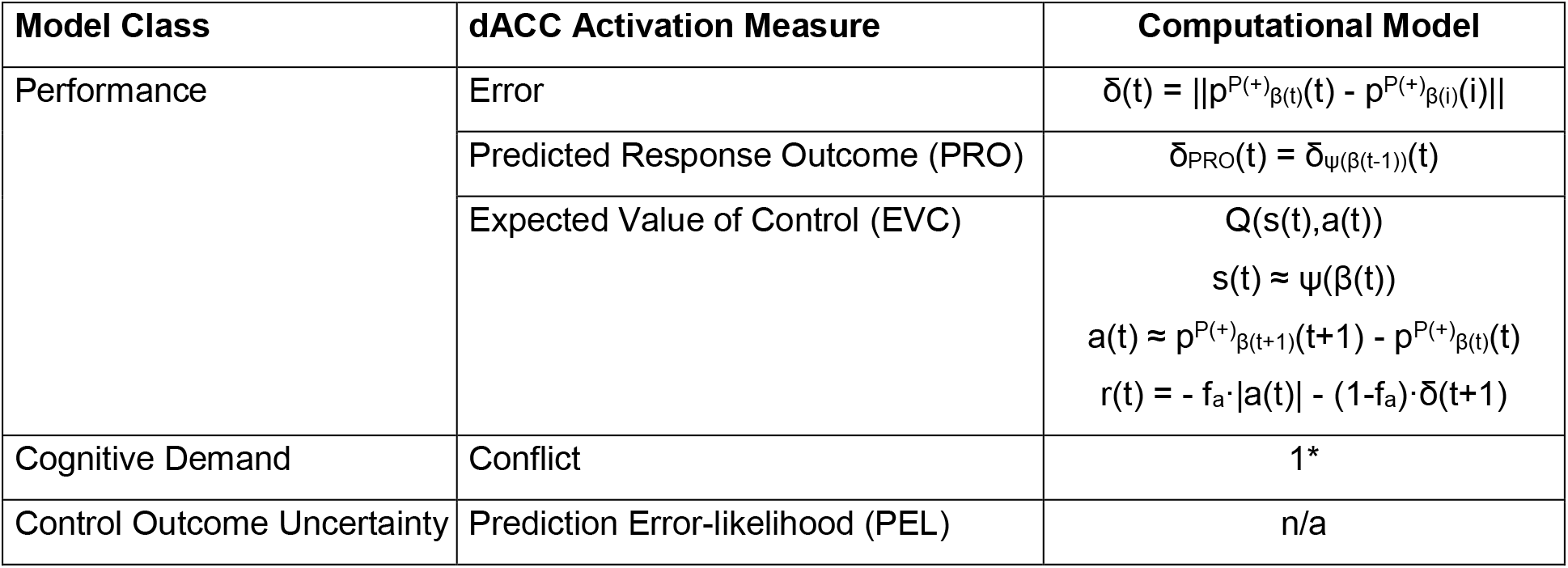
Summary of cognitive control models of dACC function. *Theorized model of conflict assuming that “non-action” is a valid prepotent response to the cued-recall/re-experiencing task.

The remaining two performance monitoring-related models, cognitive conflict^39^ and prediction error likelihood (PEL)^24^, describe important theoretical aspects of cognitive control that are related to our three included models and may be subsumed by, or derived from, them. However, PEL research has shown that the dACC activates in response to contexts in which there is a learned probability of failure, even when acting rationally. It is questionable as to whether the neural mechanisms underlying explicit affect regulation conform to such a highly stochastic transition function. Moreover, our analysis plan relied on the subject’s awareness of the affective goal state throughout the cued-recall/re-experiencing trial, thereby conceptually excluding this model from our comparative analysis. Similarly, our task design did not include a canonical test of cognitive conflict. We had two lines of reasoning that supported this research design. First, our previous work showed that affect processing was neurally encoded orthogonally across the independent affective dimensions of valence and arousal^47^; therefore, affect processing should not be dependent on a conflict signal in the dACC. Second, our most plausible assumption of the prepotent response to explicit affect regulation was that of “non-action”. Baseline dACC activation during the regulation demand of the cued-recall/re-experiencing task would suggest cognitive control recruitment to overcome this prepotent response and, therefore, conflict did not need to be modeled explicitly.

Error formed the foundation of our modeling approach. As indicated in Table 1 (and conceptualized by Fig. 1, panel B), we computed error, δ, as the difference in Platt-scaled decoded affect processing between each regulation-related volume and that related to the target cue response (i.e., the affective goal) separately for both valence and arousal. From these basic calculations, we constructed the remaining models.

Both the PRO and EVC models relied on predictions of derived quantities, which themselves are based upon a state space of neural activations. This posed a significant challenge in that task-related activations of the dACC are likely to be temporally correlated with other neural activations throughout the brain, potentially resulting in false positive model outcomes. To control for this possibility, we constructed a novel image processing pipeline to isolate our models of the performance monitoring signals (in both time and neuroanatomical space) from the dACC activations that were the subject of functional characterization.

First, we generated a mask of the entire medial frontal cortex (mFC) by inflating an existing mask^69^ by two voxels. We then used this mask to exclude mFC voxels from an established 20-component (18 usable components) partition of the brain derived from independent components analysis of the BrainMap database^70^. We then mapped all beta-series into this mFC-excluded 18-dimensional space. We denoted this transformation of the beta-series as Ψ(β) and refer to it as the restricted beta-series, which was used (as described below) to model task-related neural activity for neural processing networks that excluded the mFC.

We next fit linear support vector machine regression models of the computed error trajectories, using the restricted beta-series as input features, where the input feature is drawn from the previous time index as the error to be predicted (i.e., error forecasting, see Table 1, PRO). To predict each target subject’s future error, we fit within-subject regression models for each of the remaining set of subjects, using these subjects’ restricted beta-series and error trajectories for training. We then estimated the target subject’s PROs by ensemble averaging the remaining subjects’ model predictions, using the target subject’s restricted beta-series as input features to these models. This model-building step is critical in that potential temporal correlations between the dACC and the remainder of the brain are not available to be learned by the PRO decoding models. In order to validate this modeling approach, we measured the effect size of the ensemble average of the remaining subjects’ model predictions in approximating the target subject’s predictions. We measured this effect size using a linear mixed-effects model where random slope and intercept effects were modeled subject-wise. We estimated effect sizes separately for valence and arousal.

We showed by direct proof (see Supplemental Figure S2) that EVC is mathematically equivalent to Q-learning^71^ with a composite reward function that incorporates the cost of action in addition to the transition reward. We computed an approximate batch solution to the action-value function, Q, via fitted Q-iteration^72^ using the following constituent components: states, s, actions, a, and rewards, r (see Table 1, EVC). We modeled actions as the differences between successive affect processing predictions (i.e., forward Euler approximations of the first temporal derivative), respectively, for valence and arousal. We modeled states identically to the methods used for the PRO model (i.e., restricted beta-series). Finally, we modeled the reward function as the weighted combination of the action and the error (note, these equations assume the machine learning convention in which rewards can have either positive or negative sign).

We conducted all EVC experiments over a parameter space composed of the cross product of the discount factor, γ, sampled from the range [0,1] at intervals of 0.1 and fraction of action cost, f_a_, sampled from the range [0,1] at intervals of 0.2. We also discretized the action space to five actions, a_d_ ∈ [−2,−1,0,1,2] according to the following heuristic. We computed actions for each subject and standardized these values subject-wise. We then pooled all standardized actions and computed the group standard deviation, σ. We then assigned discrete actions according to a standard deviation-driven partition of the standardized action space of each subject such that

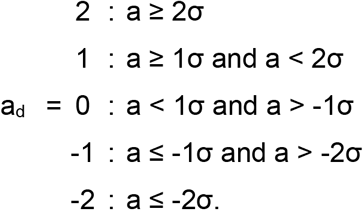

For each subject, we separately standardized the actions and errors for all cued-recall/re-experiencing task trials and then calculated the reward according to the appropriate meta-parameters for the given experiment. We then fit the action-value function according to a random forest-based implementation of fitted Q-iteration^72,73^. However, similar to our scheme for estimating PROs, we employed an out-of-sample ensemble averaging for our EVC estimate for each action at each state. We stored for additional analysis: (1) the ensemble average prediction of Q-value for the on-policy actions, (2) the expected Q-values of random action at each state (where probabilities of each discrete action were estimated from the subject’s distribution of actions), and (3) the errors between the on-policy action and the optimal action for each state.

For each set of parameters, we tested for group-level significant differences between the Q-values of on-policy actions and the expected Q-values of random actions using the Wilcoxon rank-sum test. Within identified significant parameter sets, we calculated the group-level mean error between on-policy actions and optimal actions. We then selected the meta-parameter set such that group-level mean error was minimized. We broke ties as follows: maximum Ɣ (to decorrelate EVC from PRO) and minimum f_a_ (we assumed cognitive effort was subordinate to control performance).

### Comparing Performance Monitoring Models of Medial Prefrontal Cortical Activation

We critically compared our constructed models of performance monitoring using whole-brain gray matter linear mixed-effects models, implemented via AFNI’s 3dlme function. To control for our use of neural activations outside the dACC to predict model values within the dACC, we constructed two separate mixed-effects models. The first model (see Model 1) incorporated the performance monitoring fixed effects. We used this model to characterize only neural activations within the dACC. The notation (Fixed_i_ × Sex × Age) represents an expansion of fixed effects representing all possible interactions of age and sex with each of the explicitly listed primary fixed effects, Fixed_i_, which by default also included pair-wise interactions, e.g. Fixed_i_ × Sex and Fixed_i_ × Age as well as Sex and Age alone.

***Model 1:***

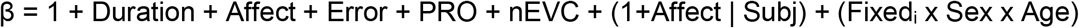

We used the second model (see Model 2), which omitted the measurements of performance monitoring as fixed effects, to characterize neural encodings falling outside of the dACC.

***Model 2:***

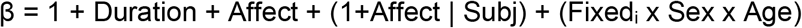

The resulting solutions provided F-statistics for each fixed-effect (and interaction) for each voxel included in the model. We also constructed general linear tests for all primary fixed effects of interest to characterize the sign of the effect sizes for subsequent analysis. An important detail, we negated our EVC predictions (denoted nEVC) to enforce semantic symmetry among all performance monitoring regressors such that positive values indicated poor quality states and negative values indicated high quality states across all three models (Error, PRO, and nEVC).

We computed cluster-size thresholds according to recently reported methodological advances^74^. First, we used the mixed-effects model residuals to estimate the shape parameters of the spatial autocorrelation function (ACF) via AFNI’s 3dFWHMx function. We then simulated, via AFNI’s 3dClustSim function (-acf option), the cluster-size threshold for our voxel-wise statistical maps (thresholded to p<0.001, uncorrected) that survive family-wise error (FWE) corrected thresholds of p < 0.05, assuming clusters were formed from contiguous (face-touching, NN=1) voxels. We performed this thresholding process separately for F- and Z-statistical maps.

Finally, we restricted reporting of our first model to only those voxels within a dACC mask (see Figure 3) constructed from the dACC-engaged component of an established 70-component independent component analysis-based partition of the BrainMap database^70^. Note, this mask was thresholded to remove several small activations or voxel clusters not belonging to the dACC proper. We restricted our reporting of the second model to all surviving voxels falling outside of our inflated mFC mask.

### Overview of Analyses

In our first experiment, we constructed and validated predictive models of affect processing according to whole-brain neural activation responses to image stimuli in order to confirm prior reports of prediction performance in this class of model. In our second experiment, we applied these models to measure trajectories of affect processing from whole-brain neural activation responses to the cued-recall/re-experiencing task of image stimuli in order to test the effect size of explicit affect regulation against affect regulation occurring at rest. In our third experiment, we converted the cued-recall/re-experiencing task’s affect processing trajectories into estimates of affect regulation-related performance monitoring relative to the task goal and critically tested their association with activations in the dACC. In our fourth experiment, we characterized the neural activations associated with the cued-recall/re-experiencing task itself to contextualize our findings within the extant emotion regulation literature.

## Results

### Neural Decoding Models Accurately Classify Implicitly Induced Affect Processing States

We constructed linear support vector machine (SVM) classifiers for 89 subjects (excluding 5 subjects for excessive head motion) based upon responses to 90 image stimuli computationally sampled from the IAPS. We observed classification accuracies (see Supplemental Table S4) that were both group-level significant (p<0.001; Wilcoxon signed rank; h_0_: μ=0) and consistent with the highest classification accuracies reported in the literature when controlled for the affect processing induction reliability of stimuli, which was previously shown to be a function of the stimuli’s affective properties^48^. We also found that 79 of 89 subjects (88.8%) exhibited significant within-subject classification of affective valence and arousal stimuli, respectively (p<0.05; binomial distribution, h_0_: within-subject permutation test accuracy). Further, we controlled for the dimensionality of the feature space^48^ by relating affective decodings predicted using whole-brain gray matter voxel-wise features (30,000–40,000 dimensions) to decodings predicted using Gram-Schmidt dimensionally-reduced features (90 dimensions, see Supplemental Figure S3), which confirmed that our decoding models scaled well to the whole-brain gray matter feature space. As further evidence of the validity of the neural decoding models, conversion of the decoding models into their equivalent encoding models via Haufe-transform^75^ confirmed that the resultant encoding models broadly recapitulate the established distribution of the neural correlates of affective valence and arousal processing identified through both univariate^76^ and multivariate^48,77,78^ methodology (see Supplemental Figure S4). These results support the ability of the IAPS stimuli to induce the expected affect processing responses and the ability of the neural decoding models to accurately classify and isolate their putative neural processing correlates. To the computational modeling approach, these results also support the assumption that the plant is observable based on the ability to measure its state, s.

In this work, we applied these neural decoding models to measure moment-to-moment affect processing states during affect regulation. Our control theoretic analysis presumed that these state measurements conform to a continuum of real-values. Therefore, we converted our predictions of hyperplane distance (i.e., classification label assignments) into probabilities via Platt scaling^67^, using the probability of affect state membership to the positive class (i.e., positive valence or high arousal) as our measurement convention. We then validated this conversion by predicting the Platt-scaled known normative scores of IAPS training stimuli from Platt-scaled within-subject neural decodings (see Supplemental Figure S5) and demonstrated significant prediction performance, respectively, for valence (p<0.001; t-test; h_0_: β=0) and arousal (p<0.001; t-test; h_0_: β=0). These results support our approach to neural pattern classification as a means of tracking momentary affect processing states and thus the ability to temporally characterize the performance monitoring (control law, L) function of affect regulation.

We also extended our validation of the decoding models to include those image stimuli reserved for the cued-recall/re-experiencing task (a distinct set of 30 image stimuli sampled independently, but identically, to the training image set). We tested the prediction accuracy of the Platt-scaled known normative affect dimension scores of these task cue image stimuli using Platt-scaled within-subject decodings (see Supplemental Figure S6) and, again, demonstrated significant predictive effects, respectively, for valence (p<0.001; t-test; h_0_: β=0) and arousal (p<0.001; t-test; h_0_: β=0).

### Independent Validation of Experimentally-Induced Affect Processing

We also employed independent psychophysiological measures of affect processing to confirm the induction of affect processing for trials on which we trained our predictive models, a critical manipulation check^79^. We predicted the known normative scores of the stimulus set from their induced facial electromyography and electrodermal responses, respectively, for the independent affective dimensions of valence and arousal. We found that corrugator supercilii electromyography is a weak (R^2^=.0007), but significant (p=.024; t-test; h_0_: *β*=0), index of IAPS stimuli valence scores (see Supplemental Figure S7 as well as Supplemental Figure S8, panel A) when applied to detect valence in polar-extreme implicit stimuli (thresholded to remove approximately the middle third of stimuli as measured by normative score). Similar analysis of zygomaticus major electromyography was not significant (p=0.78; t-test;; h0: *β*=0). We also found electrodermal activity to be a weak (R^2^=.002), but significant (p<0.0002; t-test; h_0_: *β*=0), index of IAPS stimuli arousal score (see Supplemental Figure S8, panel B), which was consistent with previous reports using similar methods^47^.

### Estimating Affect Regulation Effect Sizes within the Cued-Recall/Re-experiencing Task

We applied our models to decode moment-to-moment trajectories of affect processing under regulation during the cued-recall/re-experiencing task for (n=86) subjects (we excluded 3 additional subjects due to head motion censoring that precluded extraction of task-related neural activations via the beta-series method). We measured explicit affect regulation according to a linear mixed-effects model in which the fixed-effects were the standardized Platt-scaled predictions of affect processing-related brain states in response to the cue stimuli as well as the times elapsed since presentation of the cue. The measurements of interest were standardized Platt-scaled predictions of affect processing for the recall/re-experiencing-related fMRI volumes. We modeled random slope and intercept effects subject-wise. For each subject we also constructed a cued-recall/re-experiencing task control condition composed of neural activations extracted from 30 uniformly randomly sampled time-points within the subject’s resting state fMRI scan. We found a significant explicit affect regulation effect related to the goal state corresponding to the individual affective response properties of the cue stimulus (see Figure 2) across both affective dimensions. Further, we demonstrated that, due to its explicit and goal-directed nature, this regulation effect was significantly greater than that of the resting state task control effect for both valence and arousal.

**Figure 2:**
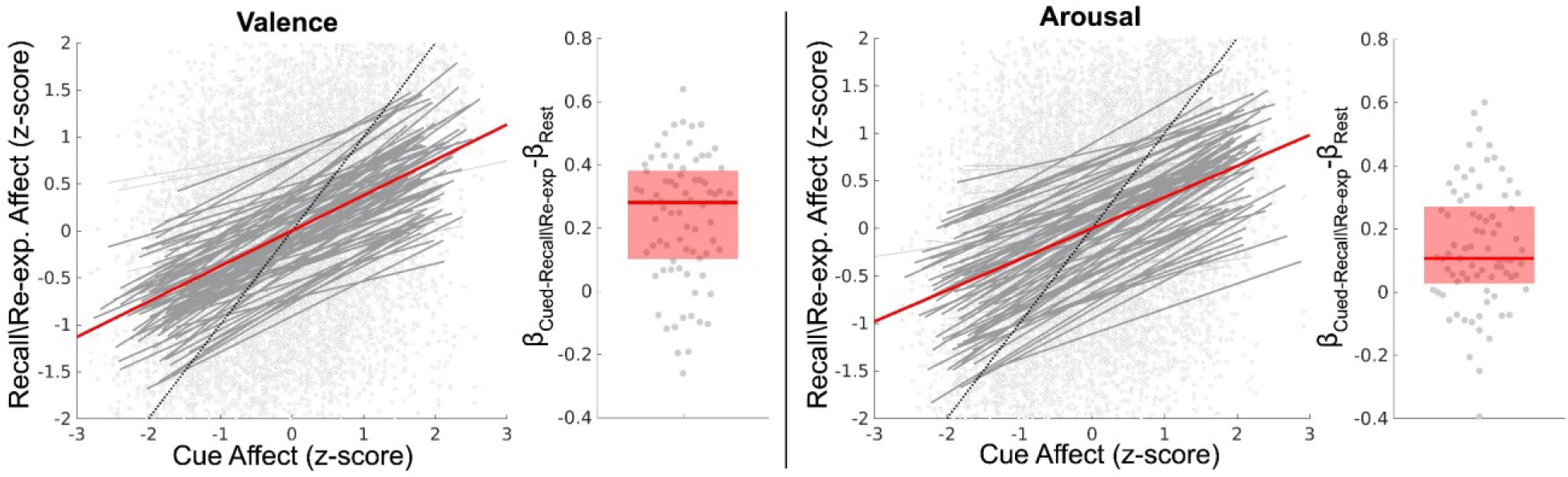
Estimation and validation of explicit affect regulation effects within the cued-recall/re-experiencing task. The figure depicts the effect size of cue affect processing in explaining the regulation of affect processing occurring during recall/re-experiencing (controlling for time lag in the 4 repeated measures of recall per each cue condition). Here affect processing measurements are standardized Platt-scaled neural decodings from our fitted SVM models. Valence and arousal dimensions of affect are decoded separately. The figure’s scatterplots depict the group-level effects computed using linear mixed-effects models which model random effects subject-wise. Bold red lines depict group-level fixed-effects of the cued affect. Bold gray lines depict significant subject-level effects whereas light gray lines depict subject-level effects that were not statistically significant. The figure’s boxplots depict the group-level difference between each subject’s explicit affect regulation measured during the cued-recall/re-experiencing task in comparison to non-explicit affect regulation constructed from the resting state task. The bold red line depicts the group median difference in effect size between cued-recall/re-experiencing and resting state. The red box depicts the 25–75th percentiles of effect size difference. **Valence**. The fixed effect (R^2^=.15) is significant (p<0.001; t-test; h_0_: β = 0). Random effects significantly improve effect-size (p<0.05; likelihood ratio test; h_0_: observed responses generated by fixed-effects only). Cued-recall/re-experiencing affect regulation effects are significantly greater than that of resting state effects (p<0.001; Wilcoxon signed rank; h_0_: β_Cued-Recall_-β_Rest_=0). The fixed-effect of control condition duration is significant (β =.02; p=.035; t-test; h_0_: β = 0). **Arousal.** The fixed effect (R^2^=.11) is significant (p<0.001, t-test; h_0_: β = 0). Random effects significantly improve effect-size (p<0.05; likelihood ratio test; h_0_: observed responses generated by fixed-effects only). Cued-recall/re-experiencing affect regulation effects are significantly greater than that of resting state effects (p<0.001; Wilcoxon signed rank; h_0_: β_Cued-Recall_-β_Rest_=0). The fixed-effect of control condition duration is not significant (β=.01; p=.26; t-test; h_0_: β = 0).

**Figure 3:**
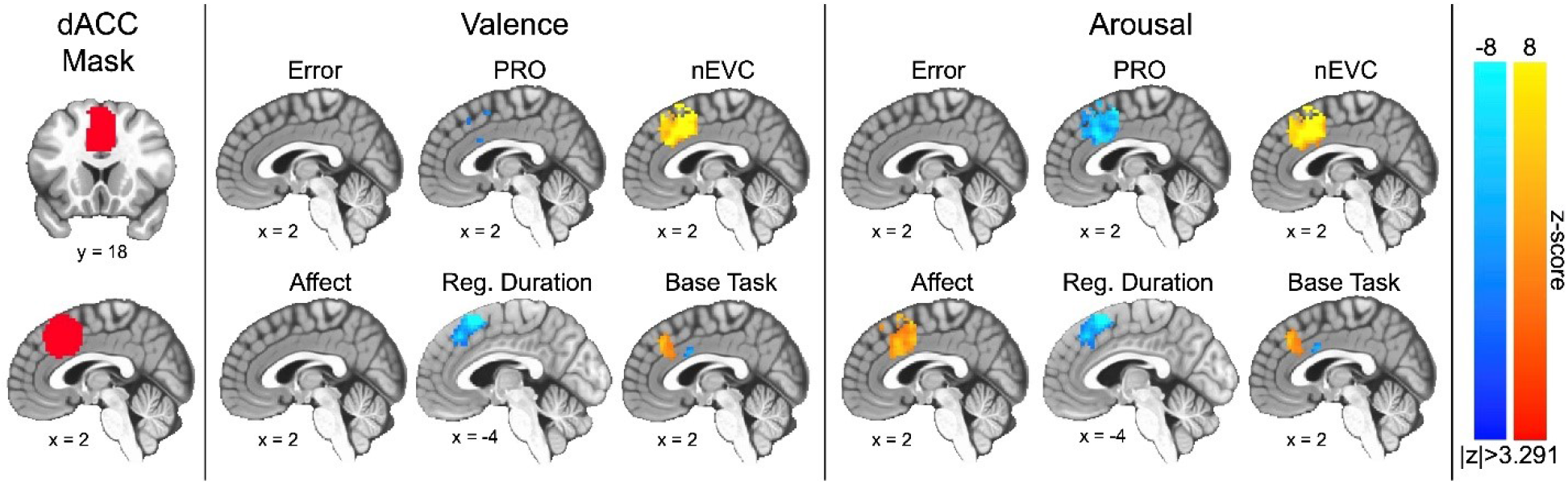
Group-level linear mixed-effect model distributions for the main fixed-effects of error, predicted response outcome (PRO), negated expected value of control (nEVC), affect processing, regulation duration, and base task (i.e., intercept) constrained to a mask of the dorsal anterior cingulate cortex (dACC). The figure depicts slices in MNI coordinate space and neurological convention (image left equals participant left). The figure depicts voxel intensities as colors having a minimum (thresholded) voxel intensity of |z| > 3.291 and a maximum absolute voxel intensity of |z| = 8.0, i.e., color saturates for absolute z-scores above the maximum intensity. To correct for multiple comparisons (p<0.05), the figure depicts only valence-derived clusters having ≥ 15.3 contiguous voxels (measured as face-wise nearest neighbors, i.e., NN=1) or arousal-derived clusters having ≥ 15.8 contiguous voxels. Note, the sagittal plane slice representing effects of regulation duration has been shifted to accommodate the entirely left-lateralized effect. Also note that an alternate sagittal slice representation of valence-derived predicted response outcome is presented in Supplemental Figure S11, which more fully captures this measure’s negative encoding.

### Independent Validation of Task-related Explicit Affect Regulation

In support of these findings of significant task-related affect regulation, we took the additional step of independently verifying, via psychophysiological measurements^79^, affect processing states in the context of explicit regulation. To this end, we constructed both the cued-recall/re-experiencing and the comparison control condition (i.e. resting state) measurements of facial electromyography (cEMG and zEMG) associated with hedonic valence^80^ as well as electrodermal activity (SCR) associated with autonomic arousal^80^. We then calculated the effects of explicit affect regulation according to linear mixed-effects modeling (see Supplemental Figure S9) and we verified the existence of explicit affect regulation-related psychophysiological responses that were positively related to the goal state psychophysiological responses induced by the cue stimuli across all three psychophysiological measurements: zEMG, cEMG, and SCR. Further, we confirmed that the explicit affect regulation effects were significantly greater than the control condition (non-explicit) regulation effects extracted from resting state measurements, again, across all three physiological measurements.

### Functional Neuroanatomical Correlates of Affect Regulation-related Performance Monitoring

Having established the neural and psychophysiological correlates of task-related explicit affect regulation we next sought to identify the most mathematically plausible theoretical model of performance monitoring encoding within the dACC. To this end, we first computed the moment-to-moment error trajectory for each cued-recall/re-experiencing trial for each subject. We partitioned our neural activation maps into two sets: activations falling within our liberal mask comprising the medial frontal cortex (mFC, including dACC), and those falling outside the mFC. We then constructed and validated out-of-sample, between-subject ensemble moment-to-moment estimates (temporally aligned with the error measurements) of both the predicted response outcome (PRO, see Supplemental Figure S10) and expected value of control (EVC, see Supplemental Table S5). We based these estimates solely on machine learning models of task-related neural activations falling outside the mFC mask. As an additional test of model performance, we computed bivariate correlation coefficients, R, between our models of performance monitoring and observed small (|R|<0.10), but significant, positive correlations between estimated expected value of control (EVC) and estimated predicted response outcome (PRO) as well as between error and PRO. We also observed small (|R|<0.10), but significant, negative correlations between EVC and error. The strength, direction, and significance of these correlations were similar across both the orthogonal affective dimensions of valence and arousal (see Supplemental Table S6).

We then estimated, via linear mixed-effects model, observed mFC neural activations as functions of error, PRO, and negated EVC (i.e., nEVC) measurements as well as affect processing measurements, time elapsed since cue onset (i.e., regulation duration), and the base task (i.e., model intercept). Note, negating the action-value fixed-effect (nEVC) enforced semantic symmetry among the three competing models of dACC activation such that poor-quality control states were associated with positive values and high-quality control states were associated with negative values. As part of this modeling process, we controlled for age and sex and included interaction terms between age and sex and all other fixed effects. Finally, we examined the specific role of the dACC in performance monitoring by restricting the model’s outputs to a dACC anatomic mask. Our findings are summarized in Figure 3.

We found that the dACC activation positively encoded performance according to the negated action-value (nEVC) as well as the base task (within the superior dACC bordering on the pre-supplementary motor area, pre-SMA). These findings were duplicated across the independent affective dimensions of valence and arousal. Concurrently, the dACC positively encoded arousal processing (see Fig. 3, Arousal: Affect). In contrast, we observed that dACC activation negatively encoded error forecasting (PRO) for both valence and arousal. Further, we observed that the left-lateralized superior dACC (bordering the pre-SMA) negatively encoded the duration of affect regulation (see Fig 3, Reg. Duration) whereas negative encoding of the base task was isolated to a small inferior and posterior region of the dACC, bordering the mid-cingulate cortex. We also observed significant dACC activation clusters supporting the interaction of age and sex effects with fixed effects in our model (see Supplemental Table S7 for a summary of these clusters).

We then modeled effects for the second neuroanatomical partition of the neural response data, those activations falling outside the mFC, using affect processing and regulation duration as well as age and sex effects and their interactions with the prescribed fixed effects. We spatially restricted our analysis to these fixed effects in order to avoid false positive associations between these data and the performance measurements that we constructed from these data using machine learning models. We summarize our findings in Figure 4.

**Figure 4:**
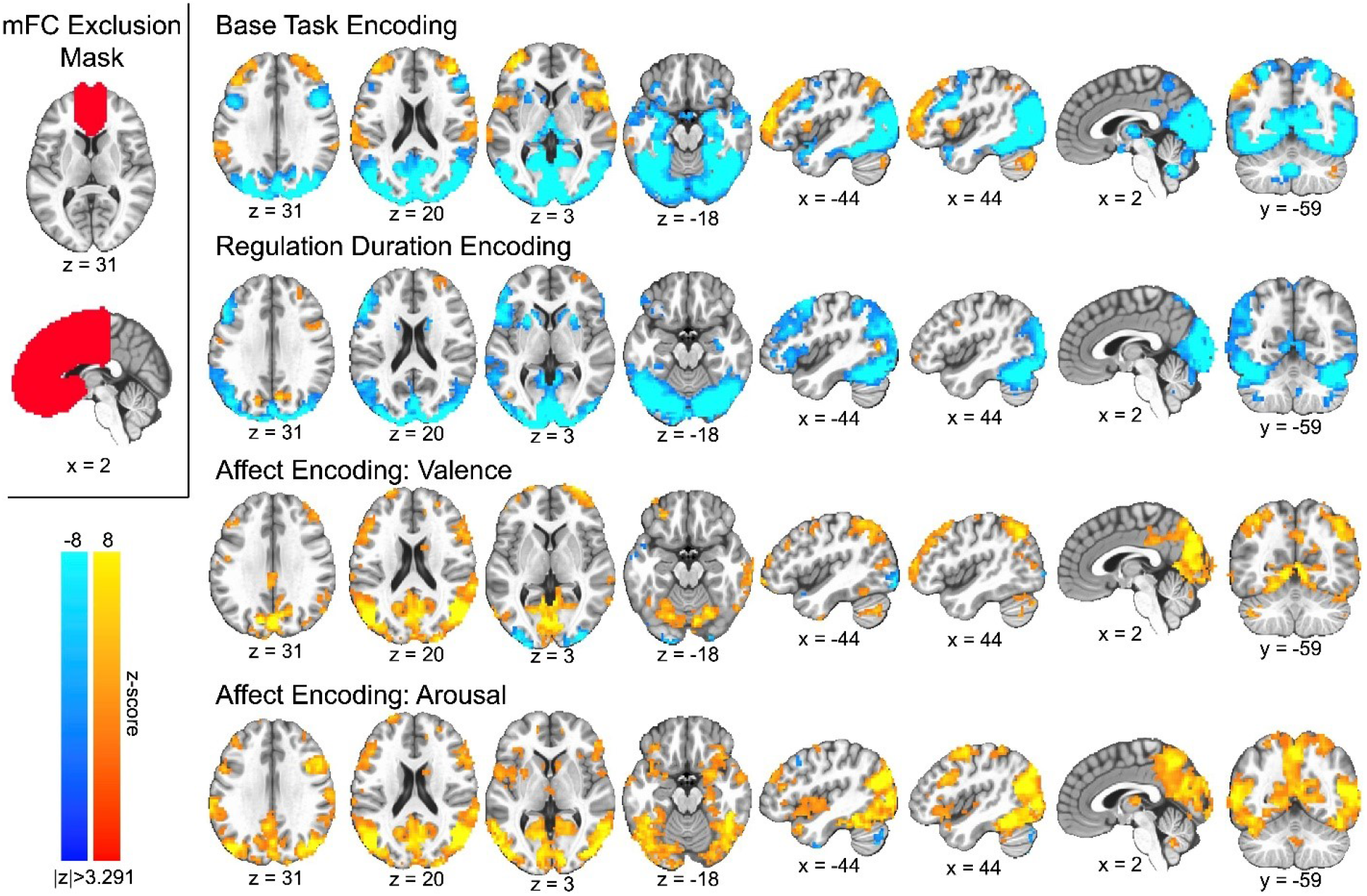
Group-level linear mixed-effect model distributions of the main fixed-effects of base task (i.e., model intercept), regulation duration, and affect processing for all gray matter voxels excluding the medial frontal cortex (upper left). Note, we report only the distributions for valence as they were highly similar to those for arousal (see OSF repository full maps). The figure depicts slices in MNI coordinate space and neurological convention (image left equals participant left). The figure depicts voxel intensities as colors having a minimum (thresholded) voxel intensity of |z| = 3.291 and a maximum absolute voxel intensity of |z| = 8.0, i.e., color saturates for z-scores above and below the maximum intensity. To correct for multiple comparisons (p<0.05), the figure depicts only valence-derived clusters having ≥ 14.9 contiguous voxels (measured as face wise nearest neighbors, i.e., NN=1) or arousal-derived clusters having ≥ 16.5 contiguous voxels.

The base task activated bilateral dmPFC, vlPFC, parietal cortex, and cerebellum as well as the left temporal parietal junction. The base task was also associated with bilateral deactivation in the striatum, dorsolateral prefrontal cortex, temporal poles, and posterior temporal cortex as well as the medial occipital cortex. We observed regulation duration to be associated with broad bilateral deactivation of the striatum, visual cortex, and ventral visual stream as well as left-lateralized frontoparietal network deactivation. Finally, we localized affect processing encodings that agreed well with distributed activations related to prior multivariate representations drawn from similar subjects in a similar task environment^47,77,81^. Again, we detected significant clusters implicating age and sex effects and their interactions with fixed effects in our model. Due to the large number of surviving clusters, we provide direct access to these cluster maps via our Open Science Framework repository (see Source Code and Data Availability).

## Discussion

The findings of this study constitute two important contributions to the affect regulation literature. First, we constructed affect processing measurement instruments, derived from machine learning predictive models, to neurally decode moment-to-moment fluctuations in affect processing during regulation. We then applied these instruments to the cued-recall/re-experiencing task and demonstrated significant explicit affect regulation effects that were concordant across two independent, objective response measurement modalities (fMRI and psychophysiology) as well as across the orthogonal affective dimensions of valence and arousal. These measurement instruments provide a novel framework to observe, analyze, and parse the functional neuroanatomical mechanisms underlying a control theoretic description of affect regulation. The second contribution of this work was our application of these novel measurement tools to construct competing computational models of performance monitoring and to critically compare each model’s ability to explain the putative role of the dACC in affect regulation.

Task-related functional neuroanatomical maps associated dACC activation with performance monitoring (Fig 3). Extended to theoretical model testing, the results most plausibly indicate that the observed dACC activation encodes the negated action-value of the control decision (nEVC model) which was conserved across both the valence and arousal dimensions of affect regulation. We interpret these findings as suggesting that dACC activation signals the expected sum of the discounted future costs of the control decision, which, according to our parameter selection process, integrates both estimated future errors and future effort in its cost expectation.

In addition to supporting the nEVC model, our findings provide evidence to reject the PRO model of the role of the dACC in performance monitoring. We observed negatively encoded prediction response outcomes related to dACC activation, which is inconsistent with the canonical definition of PRO. Moreover, we demonstrated that the PRO and nEVC regressors were weakly but positively correlated (see Supplemental Table S6); therefore, the dACC’s differential encoding of these performance models cannot be attributed to our mixed-effects modeling.

This interpretation of the study’s findings was influenced by the consideration of several alternative explanations. First, one could view our implementation of PRO as a special case of nEVC in which there was no cost of acting (f_a_=0) and future action-value was completely discounted (Ɣ=0). During a parameter search (see Supplemental Table S5), we found that on-policy control decisions input to a Q-function built using these parameters yielded action-values significantly greater than that of a random control policy. We concluded from this finding that, in general, control actions target a goal of decreased future error (as the average random action would induce zero change relative to the goal). Second, one could interpret the dACC’s negative encoding of PRO as either deactivation associated with relatively higher error or activation associated with relatively lower error. Combined, these observations suggest that the dACC positively encodes the signal for engagement of control actions, which we indirectly detected as a forecast of decreased future error.

We also observed that the dACC positively encoded the base task via the mixed-effects model’s intercept (Fig. 3). We hypothesized two complementary sources of this encoding^20^. Cognitive control theory would suggest that the observed dACC encoding represented conflict detection and processing driven by the base task’s cognitive demand, assuming that non-action was the prepotent response (see Table 1). A process model perspective of the base task would alternatively suggest that the observed dACC encoding signaled attentional deployment preceding (and in concert with) the implementation of self-control processes. Incorporating evidence from the broader view of base task-related cortical activation (Fig. 4) we observed that bilateral parietal cortex, implicated in attentional shifting^12,82^, positively encoded the base task whereas the dorsolateral prefrontal cortex, strongly implicated in cognitive emotional control^32^, negatively encoded the base task. These observations suggested that attentional deployment, rather than cognitive control, was the most plausible interpretation of the role of the dACC in the base task. Alternatively, the dACC response may reflect the role of emotion reactivity (i.e., arousal), also linked to attentional deployment^83^; however, we rejected this alternative as we explicitly modeled this possibility and observed a strong dACC encoding of affective arousal that is spatially distinct (inferior and posterior dACC).

Our analysis of the neural encodings related to the cued-recall/re-experiencing task also suggests that, compared to the canonical emotion regulation tasks of reappraisal and distraction^12^, this regulation task engages a different set of cognitive, emotional, and attentional mechanisms that may aide in disambiguating the contributions of individual neural regions to emotion regulation. Similar to reappraisal, the cued-recall/re-experiencing task exhibits activation in the vlPFC and dmPFC, suggesting a link to goal-appropriate action selection^84^ in which high-level emotional appraisal^12^ contextualizes the role of the dACC in lower level performance monitoring of the current brain state relative to a desired affective goal state. However, distinct from both the distraction and reappraisal forms of cognitive emotion regulation, cued-recall/re-experiencing did not recruit the dlPFC. This suggests that, despite this task’s explicit demand on affect regulation, it recruits an alternate mechanism relative to that for distraction or reappraisal, and, therefore, may fall elsewhere on the proposed continuum of explicit versus implicit emotion regulation tasks^33^.

### Limitations

Beyond the difficulty in disambiguating and interpreting the dACC’s encoding of prediction response outcome, our analysis excluded one important model of dACC activation (prediction-error likelihood); it also relied on a strong *a priori* assumption concerning the nature of the prepotent response to task demand in order to frame and interpret our findings within the context of cognitive conflict. We acknowledge these limitations and admit that, due to our multivariate approach to modeling dACC activation as a function of multiple performance monitoring theories, our inferences are conditional based upon the absence of explicit and convincing representations of these models. We also acknowledge that the presence of significant interactions of sex and age with several of our primary performance monitoring fixed effects suggests that the inferences we have drawn may be conditional on the demographic composition of our study’s sample. Similar analyses conducted on data from more homogeneous populations could, potentially, yield different inferences. The age and sex diversity within the studied sample, however, does provide a source of confidence concerning the generalization of inferences draw from our main fixed effects.

As with all machine learning studies, our model predictions (on which we built the primary findings of this work) relied extensively on the quality of the fMRI-derived features and their labels. We have previously reported on the limitations of exploiting IAPS normative scores as affective labels for training predictive models^47,48,60^, which also likely contributed to our observed interactions of age and sex with the neural encodings of both performance monitoring and the base task. We have also previously reported limitations related to the non-critical use of galvanic skin response as a surrogate measure of autonomic arousal^47^ and as an independent source of validation of machine learning models. We acknowledge similar limitations in our application of facial electromyography to signal stimulus-related induction of hedonic valence processing in the cued-recall/re-experiencing task.

### Broader Contributions

Beyond our exploration of the computational models underlying dACC-based performance monitoring, both the conceptual model we introduced and the methods we developed in this study may represent a novel framework for future computational approaches to the study of emotion regulation. When viewed through the lens of engineering control theory (Fig. 1, panel B), our cued-recall/re-experiencing task, which elicits a quantifiable affect regulation goal and facilitates moment-to-moment self-assessment of emotion regulation dynamics, have enabled us to:

1. clarify the differences between affect processing and affect regulation^85–87^ and, for the specific form of emotion regulation we studied, dissociate control components involved in this process;
2. formalize and study affective self-awareness, previously posited as a determining factor for successful emotion regulation^86^, enabling novel discovery in this area; and,
3. envision future control theoretic explorations of clinically relevant emotion regulation strategies, e.g. distraction and reappraisal, through formal mapping to emotional goals and goal-directed actions of relevant cognitive processes.

## Conclusion

We combined established machine learning methodology for the prediction of affect processing states from fMRI BOLD signal with independent psychophysiological measures of affect processing to validate the cued-recall/re-experiencing task as a probe and model of explicit affect regulation. We then exploited the cued-recall/re-experiencing task’s design to separate the affective goal from moment-to-moment affect regulation in order to computationally model multiple measures of performance monitoring and critically test the specific association of dACC recruitment with these model variants. We found that the dACC most plausibly computes the estimated sum of discounted future costs, including both regulation error and effort. This role was conserved across the orthogonal affective dimensions of valence and arousal. Concurrently with performance monitoring, we demonstrated that the dACC directly encodes affective arousal and also likely encodes recruitment of attention and regulation response resources.

## Acknowledgements

This study was funded by National Science Foundation grant BCS-1735820 (K.A.B). Additional personnel support was provided by National Institute on Drug Abuse grant 1T32DA022981 (C.D.K). Subject recruitment for the project was supported by the UAMS Translational Research Institute (TRI) through the National Center for Advancing Translational Sciences (1U54TR001629-01A1). The authors would like to thank Bradford S Martins, Jennifer Payne, Emily Hahn, Natalie Morris, Nathan Jones, and Laura Spell for their assistance in recruiting and assessing research subjects and acquiring subject data. The authors also thank Favrin Smith for her efforts in gaining the study’s IRB protocol approval and maintaining human subject research compliance throughout the study’s duration.

## Authorship Contributions

<u>Conception</u>: K.A.B. <u>Design, implementation, and testing</u>: K.A.B., G.A.J; <u>Analysis</u>: K.A.B, G.A.J., A.A.P, K.P.F.; <u>Interpretation of results, manuscript preparation, and revisions</u>: K.A.B, G.A.J., A.A.P, K.P.F, C.D.K.

## Competing Interests

The authors declare no competing interests.

## Source Code and Data Availability

The authors have made the source code used in this analysis publicly available: https://github.com/kabush/IN. All reported functional neuroanatomical activation maps are publicly available via the study’s Open Science Framework repository: https://osf.io/jwv6c/. De-identified CTM study data is also publicly available as a Brain Imaging Data Structure (BIDS)^88^ formatted dataset: https://openneuro.org/datasets/ds003831/versions/1.0.0. Note, this dataset does not include the (n=19) INCA study subjects who did not consent to public release of their de-identified data. For replication purposes only, de-identified raw data from the INCA study may be made privately available upon request.

## Notes

https://github.com/kabush/IN

https://osf.io/jwv6c/

https://openneuro.org/datasets/ds003831/versions/1.0.0

### Competing Interest Statement

The authors have declared no competing interest.

### Summary of Updates

MANUSCRIPT CHANGES: Replaced all references to F-statistic with t-statistic (typo). SUPPLEMENTAL MATERIALS CHANGES: Replaced all references to F-statistic with t-statistic (typo); Included Table of Contents and reformatted.

