## Supplementary Materials for "Action-value processing underlies the role of the dorsal anterior cingulate cortex in performance monitoring during self-regulation of affect"

### Supplemental Information

#### Supplemental Methods

Image Stimulus Selection
Reliable Stimulus Subset Sampling

#### Supplemental Results

| | |
|---|---|
| Supp. Figure S1. | Summary of IAPS image stimulus selection |
| Supp. Figure S2. | Proof demonstrating the equivalence of EVC and Q-learning |
| Supp. Figure S3. | Comparing decodings between GM features and Gram-Schmidt features |
| Supp. Figure S4. | Neural encodings of affect processing |
| Supp. Figure S5. | Validation of Platt-scaled decodings of affect processing |
| Supp. Figure S6. | Validation of Out-of-sample decodings of affect processing |
| Supp. Figure S7. | Facial electromyography sensitivity analysis |
| Supp. Figure S8. | Psychophysiological validation of implicit affect processing induction |
| Supp. Figure S9. | Psychophysiological validation of explicit affect regulation induction |
| Supp. Figure S10. | Validation of inter-subject ensemble PRO model of perf. monitoring. |
| Supp. Figure S11. | Detailed neural activation maps associating dACC and PRO |

| | |
|---|---|
| Supp. Table S1. | IAPS Image stimuli details |
| Supp. Table S2. | Implicit induction stimuli class counts. |
| Supp. Table S3. | Implicit induction stimuli class normative affect score distributions. |
| Supp. Table S4. | Affect processing decoding performance |
| Supp. Table S5. | Validation of inter-subject ensemble EVC model of perf. monitoring |
| Supp. Table S6. | Bivariate correlations between performance monitoring models. |
| Supp. Table S7. | Neural activation clusters of age and sex related interactions with perf. monitoring |

#### Supplemental References

### Supplemental Methods

Image Stimulus Selection

As reported in prior work[1], 90 implicit induction image stimuli were selected from the International Affective Picture Set (IAPS). Each IAPS image is associated with normative scores (based upon group-level mean measurements from a 9-point Likert scale) of image valence and arousal. We computationally sampled our image subset from the full IAPS image set according to a maximum separation heuristic in which (starting with a randomly selected image) each additional image is selected such that its normative valence and arousal scores exhibit the maximum summed Euclidean distance (in arousal-valence coordinates) to all currently selected images (see Supplemental Figure S1). 30 images comprising the cues of the cued-recall/re-experiencing trials were similarly sampled (see Supplement Figure S1). The image sets were then fixed for all participants. Normative valence and arousal scores for all image stimuli, as well as counts and distributions of the affective scores comprising the positive and negative classes used to train the decoding models, are presented in Supplemental Tables S1–S3.

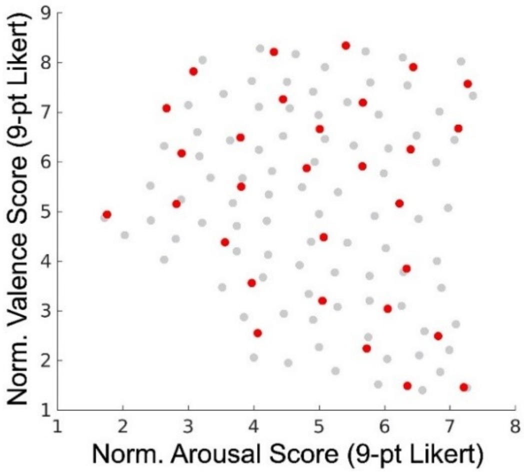


***Supplemental Figure S1:*** *Summary of the affective properties of the implicit induction and cued-recall/re-experiencing trial image stimuli based on IAPS reported normative scores. Solid red markers depict (n=30) individual cue stimuli plotted in coordinates representing mean normative arousal and valence scores. For comparison, solid gray markers depict (n=90) individual implicit induction stimuli plotted in similar coordinates.*

Reliable Stimulus Subset Sampling

A key question facing affect processing researchers is how to accurately classify stimuli. As valence and arousal are dimensional properties of affect, the natural demarcation of, e.g., positive versus negative, valence is potentially unclear *a priori*. Moreover, the propriety of labeling certain stimuli as neutrally valent is also unclear. Similar questions arise in the labeling of arousal. These questions become particularly relevant when curating stimuli for the training of affect processing decoding models and for comparing performance between decoding models. Training data that cluster in the extremes of an affective dimension may confer a degree of simplicity to the decoding problem that does not necessarily exist in nature. In prior work we described how image stimuli could exhibit reliable properties of positive versus negative affect processing by measuring the degree to which the brain states induced by these stimuli cluster. Stimuli exhibiting canonical two-class induction properties were labeled as part of the 'Reliable Stimulus Subset' of the total stimulus set[1].

In this work we modified the original Reliable Stimulus Subset selection algorithm to incorporate a null distribution formed from global permutation testing of the decoding models. For clarity, we summarize the algorithm as follows. For each subject, i, for each stimulus, j, we evaluated the reliability of stimulus j based on the distribution of predictions made for this stimulus by the remaining set of study subjects (n=88). Each stimulus that exhibited prediction accuracy greater than chance according to the binomial distribution (n=88, $\alpha$=0.05) where the null probability was defined as the mean accuracy of all remaining subjects' permutation tests (SVM fit of beta-series to uniformly randomly assigned class labels averaged over 1000 trials) was identified as 'reliable' and added to the subject's reliable stimulus set, $RSS_i$. We then conducted within-subject classification of the RSS using the $i^{th}$ subject's decoding model and report accuracy on this dataset. Reporting RSS classification accuracy, in addition to classification accuracy on the full dataset, is important for understanding potential performance biases that may exist for curated datasets in which stimuli were hand-selected and, therefore, may possess artificially clustered normative affect scores, the existence of which we have previously reported[1,2].

### Supplemental Results

$$
\begin{aligned}
EVC(s_t, a_t) &= -C(a_t) + \textstyle\sum_i p(s_i|\, s_t, a_t)(r(s_{t+1}) + \gamma \cdot max_j EVC(s_i, a_j)), from\ Equations\ 1\ \&\ 2^3 \\
&= -C(a_t) + \textstyle\sum_i p(s_i|\, s_t, a_t) \cdot r(s_{t+1}) + \textstyle\sum_i p(s_i|s_t, a)\gamma \cdot max_j EVC(s_i, a_j) \\
&= -C(a_t) + \mathbb{E}(r(s_{t+1})) + \mathbb{E}(\gamma \cdot max_j EVC(s_{t+1}, a_j)) \\
&= -C(a_t) \cdot p(1) + \mathbb{E}(r(s_{t+1})) + \mathbb{E}(\gamma \cdot max_j EVC(s_{t+1}, a_j)) \\
&= \mathbb{E}(-C(a_t)) + \mathbb{E}(r(s_{t+1})) + \mathbb{E}(\gamma \cdot max_j EVC(s_{t+1}, a_j)) \\
&= \mathbb{E}(-C(a_t) + r(s_{t+1}) + \gamma \cdot max_j EVC(s_{t+1}, a_j)) \\
&= \mathbb{E}\left(r'_{t+1} + \gamma \cdot max_j EVC(s_{t+1}, a_j)\right), where\ r'_{t+1} = \beta_1 \cdot r(s_{t+1}) - \beta_2 \cdot C(a_t) \\
&\equiv Q(s_t, a_t)\ by\ definition^4
\end{aligned}
$$

***Supplemental Figure S2:*** *Direct proof demonstrating that the expected value of control (EVC) is equivalent to a Q-function with a compound reward that incorporates the cost of action.*

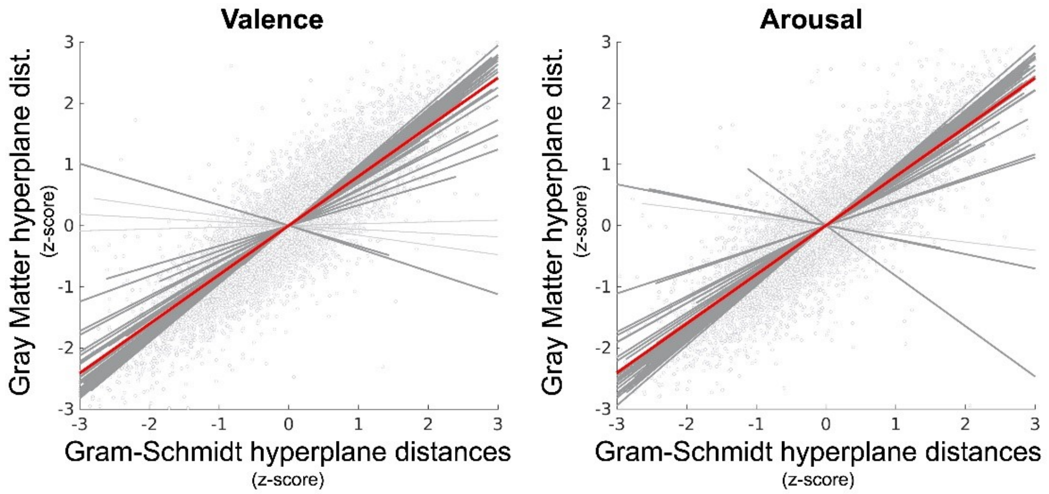


***Supplemental Figure S3:*** *Comparison of support vector machine predictions based upon whole-brain gray matter versus Gram-Schmidt reduced dimensionality features. Gram-Schmidt dimensionality reduction projects the original whole-brain gray matter features (n~30,000–40,000) onto an orthogonal basis in which the coordinate dimension is less than or equal to the number of sample features (n≤90). We report the effect size of the reduced dimensional predictions in explaining predictions in the original feature space using a linear mixed-effects model in which random effects are modeled subject-wise. Gray symbols depict individual trials. The bold red line depicts the group-level effect. Bold gray lines depict significant subject-level effects whereas light gray lines depict subject-level effects that were not significant.* ***Valence.*** *The fixed effect ($R^2$=.71) is significant ($p<0.001$; t-test; $h_0$: $\beta=0$). Random effects significantly improve effect-size ($p<0.05$; likelihood ratio test; $h_0$: observed responses generated by fixed-effects only).* ***Arousal.*** *The fixed effect ($R^2$=.72) is significant ($p<0.001$; t-test; $h_0$: $\beta=0$). Random effects significantly improve effect-size ($p<0.05$; likelihood ratio test; $h_0$: observed responses generated by fixed-effects only).*

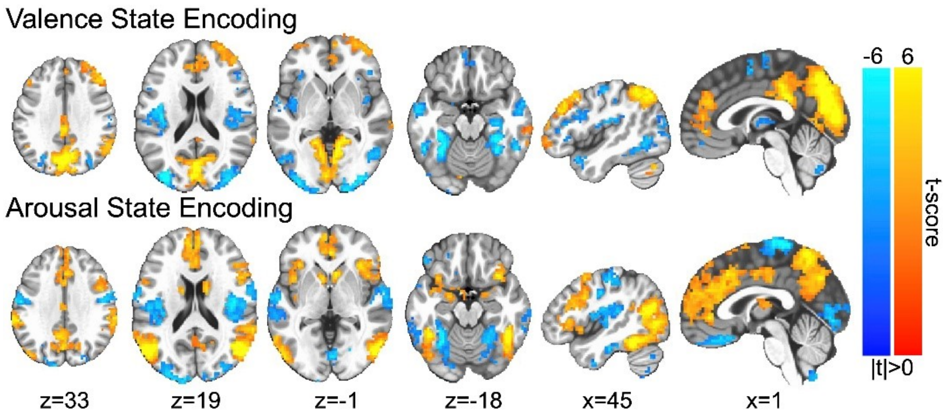


***Supplemental Figure S4.*** *Neural encodings of affect processing. Color gradations indicate the group-level t-scores of the encoding parameters (red indicating positive valence or high arousal, blue indicating negative valence or low arousal). T-scores are presented only for those voxels in which encoding parameters survived global permutation testing ($p<0.01$, uncorrected, N=1000 random permutations). Image slices are presented in MNI coordinate space and neurological convention. Maximum voxel intensity is |t|=6.0, i.e., color saturates for t-scores with absolute values falling above this value.*

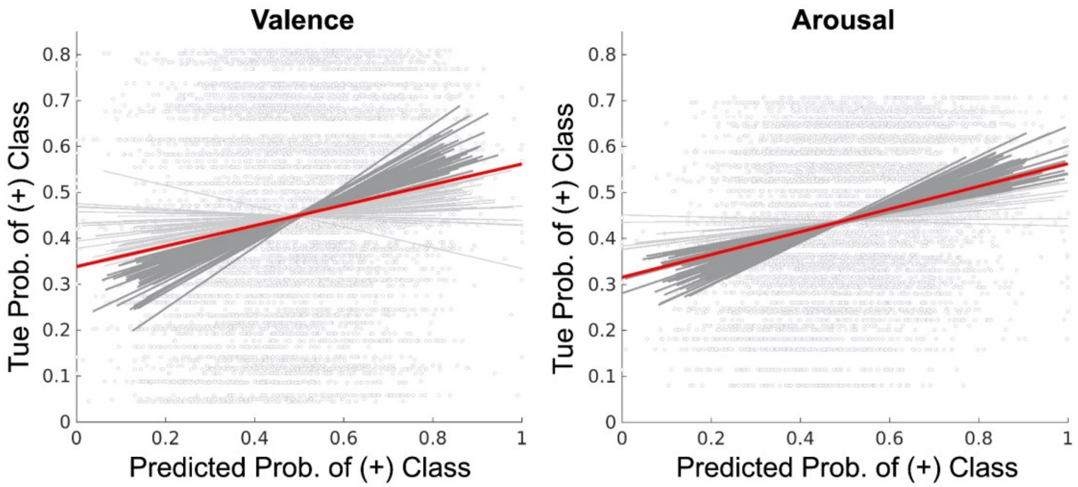


***Supplemental Figure S5:*** *Validation of Platt-scaling of the hyperplane distance predictions of affect processing. The figure depicts the effect size of Platt-scaled hyperplane distance predictions in explaining the Platt-scaled normative affect scores of IAPS stimuli used to train the support vector machine classifiers, separately for the orthogonal affective dimensions of valence and arousal. Hyperplane distance predictions resulted from within-subject leave-one-out cross-validation. The figure depicts the group-level effects computed using a linear mixed-effects model which modeled random effects subject-wise. Gray symbols depict individual trials. The bold red line depicts the group-level effect. Bold gray lines depict significant subject-level effects whereas light gray lines depict subject-level effects that were not significant.* ***Valence****. The fixed effect ($R^2$=.03) is significant ($p<0.001$; t-test; $h_0$: $\beta=0$). Random effects significantly improve effect-size ($p<0.05$; likelihood ratio test; $h_0$: observed responses generated by fixed-effects only).* ***Arousal.*** *The fixed effect ($R^2$=.07) is significant ($p<0.001$, t-test; $h_0$: $\beta=0$). Random effects do not significantly improve effect-size.*

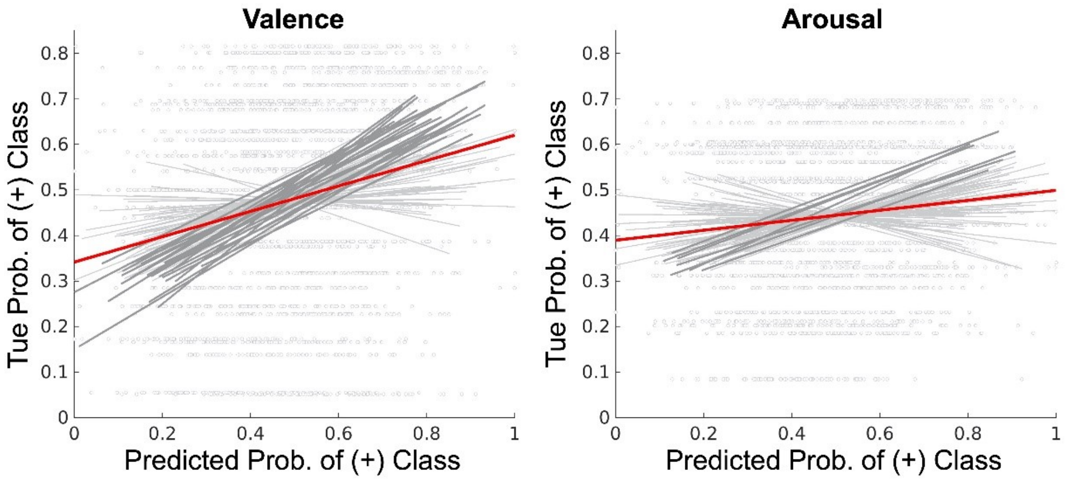


***Supplemental Figure S6:*** *Out-of-sample validation of linear support vector machine model predictions. The figure depicts the effect size of Platt-scaled hyperplane distances predicted by the fitted SVMs in explaining the Platt-scaled normative affect scores of IAPS stimuli used as cue stimuli in the cued-recall/re-experiencing affect regulation task. Effect-sizes are reported separately for the orthogonal affective dimensions of valence and arousal. The figure depicts the group-level effects computed using a linear mixed-effects model which modeled random effects subject-wise. Gray symbols depict individual trials. The bold red line depicts the group-level effect. Bold gray lines depict significant subject-level effects whereas light gray lines depict subject-level effects that were not significant.* ***Valence****. The fixed effect ($R^2$=.05) is significant ($p$<0.001; t-test; $h_0$: β=0). Random effects do not significantly improve effect size (likelihood ratio test; $h_0$: observed responses generated by fixed-effects only).* ***Arousal.*** *The fixed effect ($R^2$=.01) is significant ($p$<0.001; t-test; $h_0$: β=0). Random effects do not significantly improve effect-size (likelihood ratio test; $h_0$: observed responses generated by fixed-effects only).*

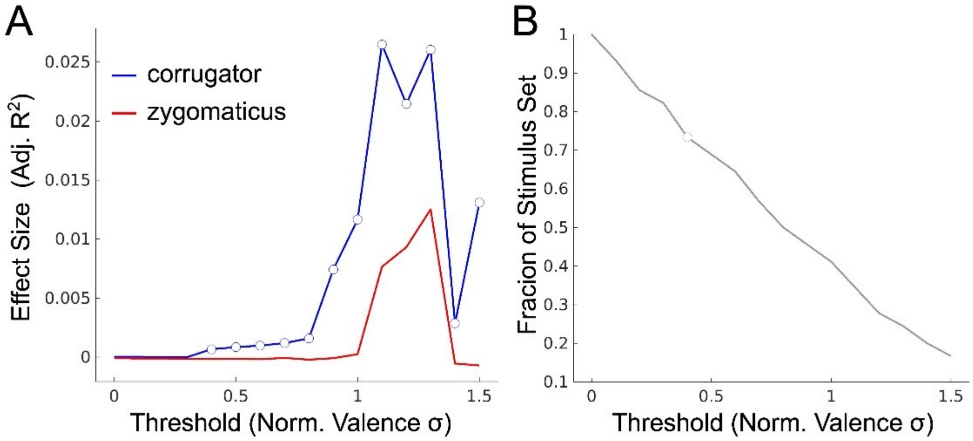


***Supplemental Figure S7:*** *Facial electromyography sensitivity analysis in the prediction of normative valence.* ***(A)*** *Valence prediction effect-size, measured as adjusted $R^2$, as a function of the polar extremes of affectively valent stimuli used to construct the prediction, plotted separately for facial EMG signals recorded from the corrugator supercilii (blue) and zygomaticus major (red). Polar-extremity is reported as a factor of the standard deviation of the normative valence scores used to threshold stimuli for exclusion from the prediction. The symbols represent thresholds for which the plotted effect-size is statistically significant.* ***(B)*** *The fraction of the total number of image stimuli remaining in the set after thresholding. The symbol denotes the minimum threshold level for which the corrugator signal significantly predicted normative valence score of the remaining stimuli.*

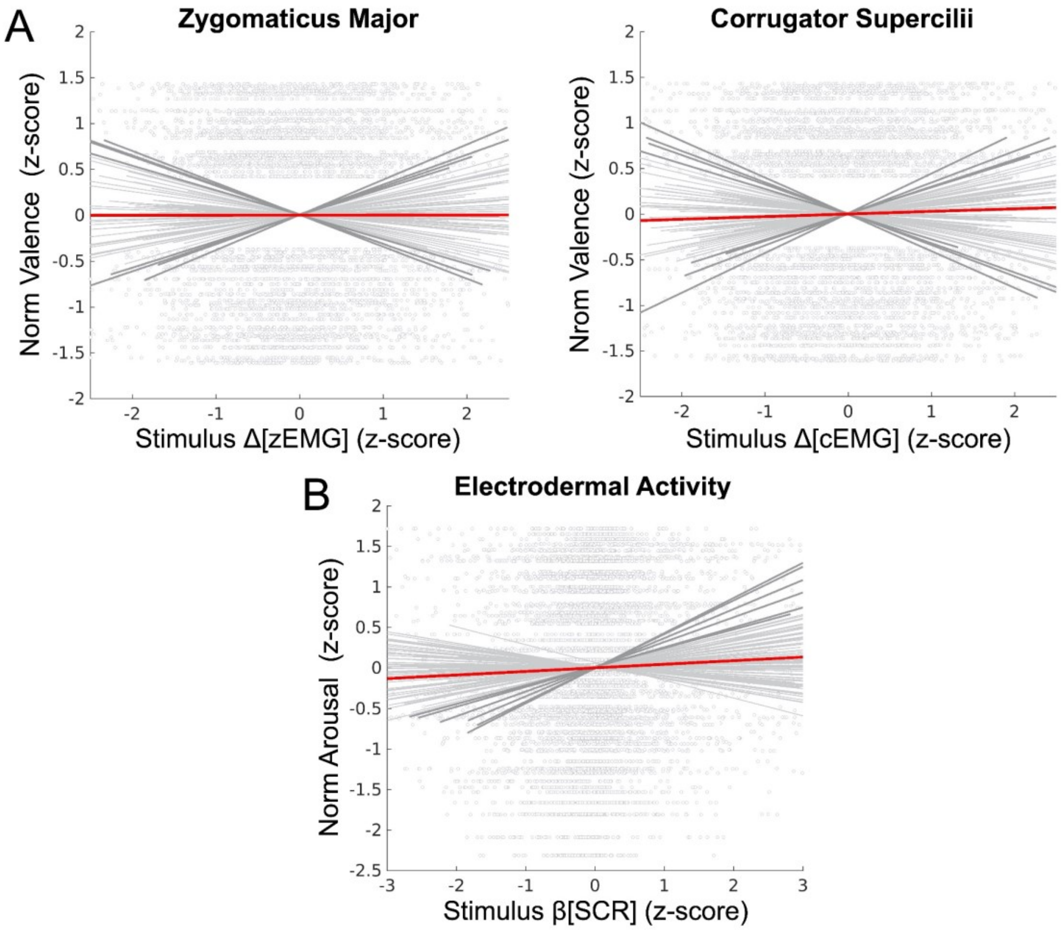


***Supplemental Figure S8:*** *Validation of psychophysiological measures as predictors of normative scores of the implicit induction stimulus set.* ***(A)*** *Facial electromyography based prediction of normative valence scores of the stimulus set (thresholded .4σ, see Supplemental Figure S7). The group-level fixed effect ($R^2$= .0002) of zygomaticus major, zEMG, differences between pre- and post-stimulus rectified signals is not significant (p=.78; t-test; $h_0$: β = 0). The group-level fixed effect ($R^2$= .0007) of corrugator supercilii, cEMG, is significant (p=.024; t-test; $h_0$: β = 0). Random effects did not significantly improve effect-size (p<0.05; likelihood ratio test; $h_0$: observed responses generated by fixed-effects only).* ***(B)*** *Electrodermal activity based prediction of normative arousal scores of the full (i.e., unthresholded) stimulus set. The group-level fixed effect ($R^2$= .002) of the skin conductance response, SCR, beta-series is significant (p<0.0001; t-test; $h_0$: β=0). Random effects did not significantly improve effect-size (p>0.05; likelihood ratio test; $h_0$: observed responses generated by fixed-effects only). In both panels, gray symbols represent individual trials, bold gray lines depict significant subject-level effects, and light gray lines depict subject-level effects that were not significant.*

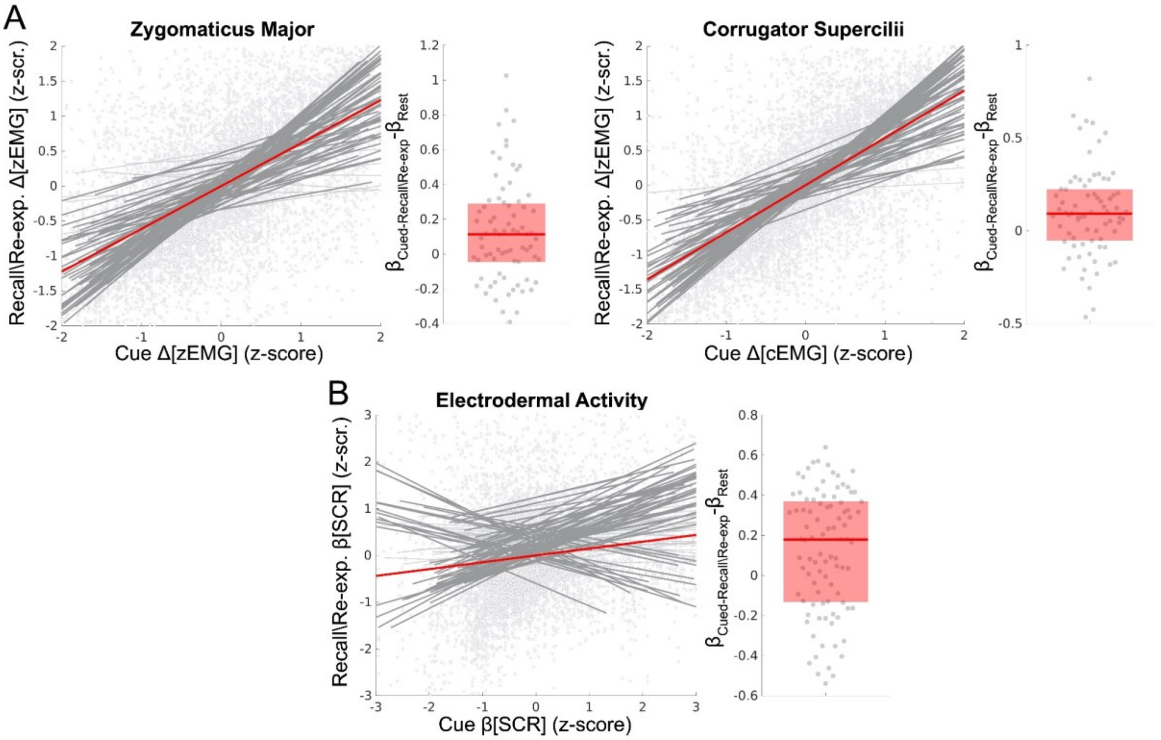


***Supplemental Figure S9:*** *Validation of explicit affect processing induction within the cued-recall/re-experiencing task via psychophysiological response measurement. The figure depicts the effect sizes of cued affect processing in explaining affect processing occurring during recall/re-experiencing (controlling for the duration of the 4 repeated measurements of recall/re-experience per each measurement of cue) for each of three unique psychophysiological measurements: facial electromyography of the zygomaticus major (zEMG), facial electromyography of the corrugator supercilii (cEMG), and electrodermal activity measured as galvanic skin conductance response (SCR). Here affect processing induction measurements are standardized measurements specific to each measurement modality (differences between pre- and post-stimulus for electromyography or modeled betas for skin conductance response). Scatterplots depict the group-level effects computed using linear mixed-effects models which model random effects subject-wise. Bold red lines depict group-level fixed-effects of the cue affect. Bold gray lines depict significant subject-level effects whereas light gray lines depict subject-level effects that were not significant. The figure's boxplots depict group-level affect processing induction measured during the cued-recall/re-experiencing task in comparison to affect processing induction constructed from the resting state task. The bold red line depicts the group median difference in effect size between cued-recall/re-experiencing and resting state. The red box depicts the 25-75th percentiles of effect size difference. Note, we measured zEMG and cEMG for CTM subjects (n=56) only. We measured SCR for all subjects.* ***(A)*** *The fixed effect ($R^2$=.45) of zEMG is significant ($p<0.001$; t-test; $h_0$: $\beta=0$). Random effects significantly improve effect-size ($p<0.05$; likelihood ratio test; $h_0$: observed responses generated by fixed-effects only). Cued-recall/re-experiencing affect processing induction effects are significantly greater than that of resting state control condition effects ($p<0.002$; Wilcoxon signed rank; $h_0$: $\beta_{IN}$- $\beta_{RST}=0$). The control duration fixed-effect is not significant ($\beta=-.006$; $p=.0415$; t-test; $h_0$: $\beta=0$). The fixed effect ($R^2$=.52) of cEMG is significant ($p<0.001$, t-test; $h_0$: $\beta=0$). Random effects significantly improve effect-size ($p<0.05$; likelihood ratio test; $h_0$: observed responses generated by fixed-effects only). The control duration fixed-effect is not significant ($\beta=-.011$; $p=.076$; t-test; $h_0$: $\beta=0$). Cued-recall/re-experiencing affect regulation effects are significantly greater than that of resting state control condition effects ($p<0.001$; Wilcoxon signed rank; $h_0$: $\beta_{IN}$- $\beta_{RST}=0$).* ***(B)*** *The fixed effect ($R^2$=.11) of SCR is significant ($p<0.001$, t-test; $h_0$: $\beta=0$). Random effects significantly improve effect-size ($p<0.05$; likelihood ratio test; $h_0$: observed responses generated by fixed-effects only). The control duration fixed-effect is significant ($\beta=-.129$; $p<.001$; t-test; $h_0$: $\beta=0$). Cued-recall/re-experiencing affect regulation effects are significantly greater than that of resting state control condition effects ($p<0.001$; Wilcoxon signed rank; $h_0$: $\beta_{IN}$-$\beta_{RST}=0$).*

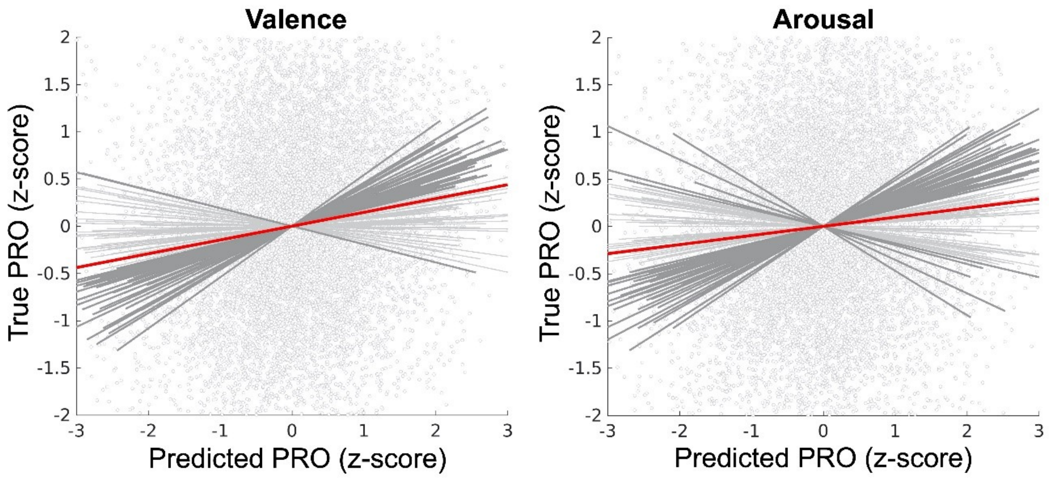


***Supplemental Figure S10:*** *Validation of out-of-sample inter-subject ensemble moment-to-moment estimates of predicted response outcome (PRO) within the dACC based upon neural activations falling outside the medial frontal cortex (mFC). Scatterplots depict the group-level effects computed using linear mixed-effects models which model random effects subject-wise. Bold red lines depict group-level fixed-effects of the models' predictions of the true PRO. Bold gray lines depict significant subject-level effects whereas light gray lines depict subject-level effects that were not significant.* ***Valence.*** *The fixed effect ($R^2$=.039) is significant ($p<0.001$; t-test; $h_0$: $\beta=0$). Random effects significantly improve effect-size ($p<0.05$; likelihood ratio test; $h_0$: observed responses generated by fixed-effects only).* ***Arousal.*** *The effect ($R^2$=.031) is significant ($p<0.001$; t-test; $h_0$: $\beta=0$). Random effects significantly improve effect-size ($p<0.05$; likelihood ratio test; $h_0$: observed responses generated by fixed-effects only).*

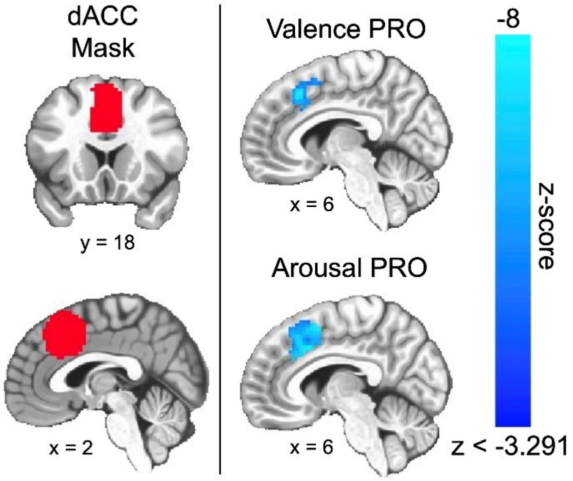


***Supplemental Figure S11:*** *Group-level linear mixed-effect model distributions for the main fixed-effect of predicted response outcome (PRO) constrained to a mask of the dorsal anterior cingulate cortex (dACC). The figure depicts slices in MNI coordinate space and neurological convention (image left equals participant left) that highlight the strongest effects of PRO (compare to Figure 3). The figure depicts voxel intensities as colors -8 < z< -3.291. Color saturates for z-scores below minimum intensity and no color is presented for z-scores above -3.291. The figure depicts only valence derived clusters having ≥ 15.3 contiguous voxels (measured as face wise nearest neighbors, i.e., NN=1) or arousal derived clusters having ≥ 15.8 contiguous voxels.*

| # | Trial Type | IAPS ID | Val. | Aro. | | # | Trial Type | IAPS ID | Val. | Aro. |
|---|---|---|---|---|---|---|---|---|---|---|
| 1 | Implicit Induction | 1022 | 4.26 | 6.02 | | 61 | Implicit Induction | 7211 | 4.81 | 4.2 |
| 2 | Implicit Induction | 1050 | 3.46 | 6.87 | | 62 | Implicit Induction | 7217 | 4.82 | 2.43 |
| 3 | Implicit Induction | 1301 | 3.7 | 5.77 | | 63 | Implicit Induction | 7224 | 4.45 | 2.81 |
| 4 | Implicit Induction | 1333 | 6.11 | 3.17 | | 64 | Implicit Induction | 7285 | 5.67 | 3.83 |
| 5 | Implicit Induction | 1620 | 7.37 | 3.54 | | 65 | Implicit Induction | 7480 | 7.08 | 4.55 |
| 6 | Implicit Induction | 1750 | 8.28 | 4.1 | | 66 | Implicit Induction | 7490 | 5.52 | 2.42 |
| 7 | Implicit Induction | 1810 | 6.52 | 4.45 | | 67 | Implicit Induction | 7492 | 7.41 | 4.91 |
| 8 | Implicit Induction | 1931 | 4 | 6.8 | | 68 | Implicit Induction | 8030 | 7.33 | 7.35 |
| 9 | Implicit Induction | 2020 | 5.68 | 3.34 | | 69 | Implicit Induction | 8158 | 6.53 | 6.49 |
| 10 | Implicit Induction | 2040 | 8.17 | 4.64 | | 70 | Implicit Induction | 8160 | 5.07 | 6.97 |
| 11 | Implicit Induction | 2058 | 7.91 | 5.09 | | 71 | Implicit Induction | 8186 | 7.01 | 6.84 |
| 12 | Implicit Induction | 2095 | 1.79 | 5.25 | | 72 | Implicit Induction | 8190 | 8.1 | 6.28 |
| 13 | Implicit Induction | 2205 | 1.95 | 4.53 | | 73 | Implicit Induction | 8200 | 7.54 | 6.35 |
| 14 | Implicit Induction | 2217 | 6.24 | 4.08 | | 74 | Implicit Induction | 8231 | 3.77 | 5.24 |
| 15 | Implicit Induction | 2222 | 7.11 | 4.08 | | 75 | Implicit Induction | 8475 | 4.85 | 6.52 |
| 16 | Implicit Induction | 2271 | 4.2 | 3.74 | | 76 | Implicit Induction | 9102 | 3.34 | 4.84 |
| 17 | Implicit Induction | 2279 | 4.71 | 3.74 | | 77 | Implicit Induction | 9120 | 3.2 | 5.77 |
| 18 | Implicit Induction | 2302 | 6.43 | 3.64 | | 78 | Implicit Induction | 9163 | 2.1 | 6.53 |
| 19 | Implicit Induction | 2351 | 5.49 | 4.74 | | 79 | Implicit Induction | 9184 | 2.47 | 5.75 |
| 20 | Implicit Induction | 2352 | 6.94 | 4.99 | | 80 | Implicit Induction | 9220 | 2.06 | 4 |
| 21 | Implicit Induction | 2520 | 4.13 | 4.22 | | 81 | Implicit Induction | 9254 | 2.03 | 6.04 |
| 22 | Implicit Induction | 2540 | 7.63 | 3.97 | | 82 | Implicit Induction | 9331 | 2.87 | 3.85 |
| 23 | Implicit Induction | 2722 | 3.47 | 3.52 | | 83 | Implicit Induction | 9360 | 4.03 | 2.63 |
| 24 | Implicit Induction | 2795 | 3.92 | 4.7 | | 84 | Implicit Induction | 9390 | 3.67 | 4.14 |
| 25 | Implicit Induction | 3000 | 1.45 | 7.26 | | 85 | Implicit Induction | 9415 | 2.82 | 4.91 |
| 26 | Implicit Induction | 3015 | 1.52 | 5.9 | | 86 | Implicit Induction | 9426 | 3.08 | 5.28 |
| 27 | Implicit Induction | 3102 | 1.4 | 6.58 | | 87 | Implicit Induction | 9435 | 2.27 | 5 |
| 28 | Implicit Induction | 3250 | 3.78 | 6.29 | | 88 | Implicit Induction | 9622 | 3.1 | 6.26 |
| 29 | Implicit Induction | 3310 | 4.37 | 5.43 | | 89 | Implicit Induction | 9700 | 4.77 | 3.21 |
| 30 | Implicit Induction | 3500 | 2.21 | 6.99 | | 90 | Implicit Induction | 9832 | 2.94 | 4.46 |
| 31 | Implicit Induction | 4220 | 6.6 | 5.18 | | 1 | Cued-Recall/Re-exp. | 1460 | 8.21 | 4.31 |
| 32 | Implicit Induction | 4235 | 5.39 | 5.29 | | 2 | Cued-Recall/Re-exp. | 1610 | 7.82 | 3.08 |
| 33 | Implicit Induction | 4490 | 6.27 | 6.06 | | 3 | Cued-Recall/Re-exp. | 1630 | 7.26 | 4.45 |
| 34 | Implicit Induction | 4503 | 6 | 4.93 | | 4 | Cued-Recall/Re-exp. | 1710 | 8.34 | 5.41 |
| 35 | Implicit Induction | 4531 | 5.81 | 4.28 | | 5 | Cued-Recall/Re-exp. | 2060 | 6.49 | 3.8 |
| 36 | Implicit Induction | 4550 | 4.95 | 5 | | 6 | Cued-Recall/Re-exp. | 2210 | 4.38 | 3.56 |
| 37 | Implicit Induction | 4597 | 6.95 | 5.91 | | 7 | Cued-Recall/Re-exp. | 2320 | 6.17 | 2.9 |
| 38 | Implicit Induction | 4598 | 6.33 | 5.53 | | 8 | Cued-Recall/Re-exp. | 3063 | 1.49 | 6.35 |
| 39 | Implicit Induction | 4619 | 6.46 | 5.09 | | 9 | Cued-Recall/Re-exp. | 3170 | 1.46 | 7.21 |
| 40 | Implicit Induction | 4626 | 7.6 | 5.78 | | 10 | Cued-Recall/Re-exp. | 4008 | 5.91 | 5.66 |
| 41 | Implicit Induction | 4641 | 7.2 | 5.43 | | 11 | Cued-Recall/Re-exp. | 4470 | 5.87 | 4.81 |
| 42 | Implicit Induction | 4649 | 5.77 | 5.99 | | 12 | Cued-Recall/Re-exp. | 4668 | 6.67 | 7.13 |
| 43 | Implicit Induction | 4770 | 4.91 | 5.85 | | 13 | Cued-Recall/Re-exp. | 5000 | 7.08 | 2.67 |
| 44 | Implicit Induction | 4800 | 6.44 | 7.07 | | 14 | Cued-Recall/Re-exp. | 5510 | 5.15 | 2.82 |
| 45 | Implicit Induction | 5010 | 7.14 | 3 | | 15 | Cued-Recall/Re-exp. | 5623 | 7.19 | 5.67 |
| 46 | Implicit Induction | 5020 | 6.32 | 2.63 | | 16 | Cued-Recall/Re-exp. | 5920 | 5.16 | 6.23 |
| 47 | Implicit Induction | 5395 | 5.34 | 4.23 | | 17 | Cued-Recall/Re-exp. | 5972 | 3.85 | 6.34 |
| 48 | Implicit Induction | 5750 | 6.6 | 3.14 | | 18 | Cued-Recall/Re-exp. | 6231 | 2.49 | 6.82 |
| 49 | Implicit Induction | 5760 | 8.05 | 3.22 | | 19 | Cued-Recall/Re-exp. | 7010 | 4.94 | 1.76 |
| 50 | Implicit Induction | 5833 | 8.22 | 5.71 | | 20 | Cued-Recall/Re-exp. | 7283 | 5.5 | 3.81 |
| 51 | Implicit Induction | 5950 | 5.99 | 6.79 | | 21 | Cued-Recall/Re-exp. | 8033 | 6.66 | 5.01 |
| 52 | Implicit Induction | 5982 | 7.61 | 4.51 | | 22 | Cued-Recall/Re-exp. | 8185 | 7.57 | 7.27 |
| 53 | Implicit Induction | 6300 | 2.59 | 6.61 | | 23 | Cued-Recall/Re-exp. | 8341 | 6.25 | 6.4 |
| 54 | Implicit Induction | 6550 | 2.73 | 7.09 | | 24 | Cued-Recall/Re-exp. | 8501 | 7.91 | 6.44 |
| 55 | Implicit Induction | 6563 | 1.77 | 6.85 | | 25 | Cued-Recall/Re-exp. | 9000 | 2.55 | 4.06 |
| 56 | Implicit Induction | 6930 | 4.39 | 4.88 | | 26 | Cued-Recall/Re-exp. | 9090 | 3.56 | 3.97 |
| 57 | Implicit Induction | 7031 | 4.52 | 2.03 | | 27 | Cued-Recall/Re-exp. | 9145 | 3.2 | 5.05 |
| 58 | Implicit Induction | 7043 | 5.17 | 3.68 | | 28 | Cued-Recall/Re-exp. | 9322 | 2.24 | 5.73 |
| 59 | Implicit Induction | 7100 | 5.24 | 2.89 | | 29 | Cued-Recall/Re-exp. | 9402 | 4.48 | 5.07 |
| 60 | Implicit Induction | 7175 | 4.87 | 1.72 | | 30 | Cued-Recall/Re-exp. | 9623 | 3.04 | 6.05 |

***Supplemental Table S1:*** *Image stimuli IAPS identifiers and normative valence (Val.) and arousal (Aro.) scores separated by trial type.*

| Class | Valence n= | Arousal n= |
|---|---|---|
| + | 45 | 44 |
| - | 45 | 46 |

***Supplemental Table S2:*** *Implicit induction stimuli class counts.*

| Class | Valence mean (s.d.) | Arousal mean (s.d.) |
|---|---|---|
| + | 6.66 (.91) | 6.09 (.68) |
| - | 3.39 (1.10) | 3.82 (.83) |

***Supplemental Table S3:*** *Implicit induction stimuli class normative affect score distributions.*

| Set of IAPS Stimuli Predicted | Valence Grp Avg. Acc. (95% CI) | Arousal Grp Avg. Acc. (95% CI) |
|---|---|---|
| Full Stim. Set (FSS) | .56 (.54,.57) | .59 (.58,.61) |
| Reliable Stim. Subset (RSS) | .77 (.75,.79) | .76 (.74,.77) |

***Supplemental Table S4:*** *SVM prediction performance.*

**Valence**

Q Performance

| | *Fraction of Action* | | |
|---|---|---|---|
| γ | 0 | 0.2 | 0.4 |
| 0 | .0389 | .1438 | .2303 |
| 0.1 | .0374 | .1380 | .2178 |
| 0.2 | .0376 | .1333 | .2107 |
| 0.3 | .0359 | .1215 | .1975 |
| 0.4 | .0344 | .1154 | .1845 |
| 0.5 | .0283 | .1116 | .1720 |
| 0.6 | .0313 | .0994 | .1600 |
| 0.7 | .0270 | .0928 | .1459 |
| 0.8 | .0248 | .0842 | .1329 |
| 0.9 | .0223 | .0786 | .1190 |
| 1.0 | .0221 | .0717 | .1084 |

Policy Error

| | *Fraction of Action* | | |
|---|---|---|---|
| γ | 0 | 0.2 | 0.4 |
| 0 | .3516 | .3293 | 0.3293 |
| 0.1 | .3451 | .3293 | 0.3293 |
| 0.2 | .3482 | .3293 | 0.3293 |
| 0.3 | .3463 | .3293 | 0.3293 |
| 0.4 | .3405 | .3293 | 0.3293 |
| 0.5 | .3428 | .3293 | 0.3293 |
| 0.6 | .3488 | .3293 | 0.3293 |
| 0.7 | .3674 | .3293 | 0.3293 |
| 0.8 | .3852 | .3295 | 0.3293 |
| 0.9 | .4920 | .3293 | 0.3293 |
| 1.0 | .4835 | .3338 | 0.3293 |

**Arousal**

Q Performance

| | *Fraction of Action* | | |
|---|---|---|---|
| γ | 0 | 0.2 | 0.4 |
| 0 | .0436 | .1457 | .2350 |
| 0.1 | .0420 | .1440 | .2275 |
| 0.2 | .0396 | .1375 | .2153 |
| 0.3 | .0400 | .1310 | .2039 |
| 0.4 | .0400 | .1236 | .1937 |
| 0.5 | .0379 | .1151 | .1801 |
| 0.6 | .0380 | .1090 | .1699 |
| 0.7 | .0380 | .1025 | .1580 |
| 0.8 | .0332 | .0957 | .1434 |
| 0.9 | .0330 | .0875 | .1343 |
| 1.0 | .0320 | .0792 | .1229 |

Policy Error

| | *Fraction of Action* | | |
|---|---|---|---|
| γ | 0 | 0.2 | 0.4 |
| 0 | .3409 | .3395 | .3395 |
| 0.1 | .3401 | .3395 | .3395 |
| 0.2 | .3407 | .3395 | .3395 |
| 0.3 | .3408 | .3395 | .3395 |
| 0.4 | .3409 | .3395 | .3395 |
| 0.5 | .3425 | .3395 | .3395 |
| 0.6 | .3419 | .3395 | .3395 |
| 0.7 | .3471 | .3395 | .3395 |
| 0.8 | .3463 | .3395 | .3395 |
| 0.9 | .3818 | .3395 | .3395 |
| 1.0 | .3828 | .3395 | .3395 |

***Supplemental Table S5:*** *Validation of inter-subject ensemble moment-to-moment estimates of expected value of control (EVC) within the dACC based upon neural activations falling outside the medial frontal cortex (mFC) and selection of optimal EVC parameters. Q Performance depicts the median action-value advantage of on-policy control versus a random policy. Policy Error depicts median squared error between the on-policy action and the optimal action. Gray cells depict the cells selected as the parameters for this experiment (see Main Manuscript Methods: Control Performance Evaluation Monitoring). Note, all parameter combinations in the Q Performance represent significant action-value advantages for on-policy control ($p<0.05$; Wilcoxon rank-sum test; $h_0$: $\mu_1$-$\mu_2$=0).* ***Valence.*** *Selected parameters: discount factor, $\gamma=0.9$; fraction of action, $f_a=0.2$.* ***Arousal.*** *Selected parameters: discount factor, $\gamma=1.0$; fraction of action, $f_a=0.2$.*

**Valence**

Correlations (R)

| | nEVC | PRO | Error |
|---|---|---|---|
| nEVC | | .0363*** | -.0506*** |
| PRO | | | .0575*** |
| Error | | | |

**Arousal**

Correlations (R)

| | nEVC | PRO | Error |
|---|---|---|---|
| nEVC | | .0552*** | -.0864*** |
| PRO | | | .0207* |
| Error | | | |

***Supplemental Table S6:*** *Summary of bivariate correlation coefficients, R, calculated between each of the primary control performance evaluation models compared in this study. *p<0.05, **p<0.01, ***p<0.001*

| Fixed-effect | # of voxels | Coordinates (CoM) x | y | z | Peak F-stat |
|---|---|---|---|---|---|
| ***Valence*** | | | | | |
| Affect x Sex | 169 | -1.2 | 16.8 | 44.7 | 40.5 |
| nEVC x Age | 26 | 3.0 | 15.6 | 29.7 | 61.2 |
| nEVC x Sex x Age | 98 | 0.5 | 12.7 | 43.3 | 52.2 |
| PRO x Age | 305 | 2.5 | 16.7 | 40.7 | 95.1 |
| PRO x Sex x Age | 77 | -4.1 | 25.0 | 37.1 | 55.8 |
| PRO x Sex x Age | 23 | 9.2 | 20.6 | 43.0 | 54.7 |
| PRO x Sex x Age | 18 | -4.9 | 24.0 | 55.9 | 34.0 |
| ***Arousal*** | | | | | |
| Affect x Age | 546 | .6 | 19.9 | 42.6 | 83.1 |
| Affect x Age x Sex | 22 | 8.7 | 23.2 | 42.5 | 25.7 |
| Error x Age | 21 | 8.2 | 22.1 | 42.9 | 30.5 |
| Error x Age x Sex | 35 | 8.2 | 23.3 | 41.7 | 22.1 |
| PRO x Age | 395 | -0.6 | 17.6 | 43.9 | 88.4 |
| PRO x Sex | 47 | -6.7 | 20.1 | 46.3 | 26.9 |
| PRO x Sex | 32 | 4.6 | 25.6 | 56.1 | 46.4 |
| PRO x Sex x Age | 166 | -3.7 | 21.4 | 42 | 56.2 |
| PRO x Sex x Age | 33 | 4.6 | 26.8 | 55.7 | 85.3 |

***Supplemental Table S7:*** *Clusters of age and sex related interactions with the performance monitoring model fixed effects separated by affect property. CoM: Center of Mass. Direct access to these cluster maps is available via our Open Science Framework repository (see Main Manuscript: Source Code and Data Availability).*
